# *foxg1a* is required for hair cell development and regeneration in the zebrafish lateral line

**DOI:** 10.1101/2024.04.12.589268

**Authors:** Jon M Bell, Cole Biesemeyer, Emily M Turner, Maddie M Vanderbeck, Hillary F McGraw

## Abstract

Mechanosensory hair cells located in the inner ear mediate the sensations of hearing and balance. If damaged, mammalian inner ear hair cells are unable to regenerate, resulting in permanent sensory deficits. Aquatic vertebrates like zebrafish *(Danio rerio)* have a specialized class of mechanosensory hair cells found in the lateral line system, allowing them to sense changes in water current. Unlike mammalian inner ear hair cells, lateral line hair cells can robustly regenerate following damage. In mammalian models, the transcription factor Foxg1 functions to promote normal development of the inner ear. Foxg1a is expressed in lateral line sensory organs in zebrafish larvae, but its function during lateral line development and regeneration has not been investigated. We find that loss of Foxg1a function results in reduced hair cell development and regeneration, as well as decreased cellular proliferation in the lateral line system. These data suggest that Foxg1 may be a valuable target for investigation of clinical hair cell regeneration.

**Summary statement:** Our work demonstrates a role for Foxg1a in developing and regenerating new sensory cells through proliferation.

## Introduction

Auditory perception and balance depend on specialized mechanosensory hair cells in the inner ear (Caprara & Peng, 2022; Fettiplace, 2017; Thomas, Cruz, Hailey, & Raible, 2015). Damage to these hair cells can occur through multiple mechanisms including genetic mutations, age, prolonged sound exposure, infection, and exposure to ototoxic drugs (Matsui & Cotanche, 2004), resulting in deafness and loss of vestibular function. Mammals are incapable of regenerating lost or damaged hair cells after development except for small populations of vestibular hair cells (Burns & Stone, 2017). This leads to permanent sensory-motor disability (Matsui & Cotanche, 2004; Roberson & Rubel, 1994). The World Health Organization estimates that as of 2018 hearing loss was the 4^th^ leading cause of disability in humans with more than 466 million individuals diagnosed (World Health Organization, 2018). In turn, the global cost of untreated hearing loss reported by the World Health Organization in 2017 was estimated to be greater than 750 billion dollars globally (World Health Organization, 2017). Research into the development of these mechanosensory hair cells and surrounding tissue is an important field for human health and economic stability.

Contrary to their mammalian counterparts, non-mammalian vertebrates are capable of regenerating functional hair cells during development and throughout adulthood (Brignull, Raible, & Stone, 2009; Ghysen & Dambly-Chaudiere, 2007; Kniss, Jiang, & Piotrowski, 2016; Pinto-Teixeira et al., 2015). In addition to inner ear hair cells, zebrafish, as well as other aquatic vertebrates, have a lateral line mechanosensory system that uses hair cells to sense water motion (Brignull et al., 2009; Coombs, Bleckmann, Fay, & Popper, 2014). Zebrafish lateral line hair cells show developmental, morphological, and genetic conservation with mammalian inner ear hair cells making them an attractive model for studying human development and disease (Nicolson, 2005). The zebrafish lateral line mechanosensory system lends itself particularly well to experimentation due to its superficial localization on the surface of the body, allowing easy visualization and manipulation. The lateral line system is made up of small sensory organs, called neuromasts, arrayed on the surface of the fish. These neuromasts contain mechanosensory hair cells and surrounding support cell (Thomas, Cruz, Hailey, & Raible, 2015). Zebrafish lateral line hair cells show dose-dependent damage in response to ototoxic drugs as is also observed in mammalian models and human patients (Brignull et al., 2009; Namdaran, Reinhart, Owens, Raible, & Rubel, 2012). Many key molecular pathways, such as FGF, Notch, and Wnt are critical for the development of inner ear and zebrafish lateral line hair cells (Baek et al., 2022; Jiang, Romero-Carvajal, Haug, Seidel, & Piotrowski, 2014; Lush et al., 2019). Support cells help form the surrounding tissue in which hair cells reside, provide trophic support for innervating neurons, and importantly for our work, act as a pool of progenitor cells for hair cell regeneration (Cruz et al., 2015; Thomas et al., 2015). Support cells’ ability to act as a progenitor pool is also regulated by FGF, Notch, and Wnt signaling (Lush et al., 2019; Lush & Piotrowski, 2014; Megerson et al., 2024). Taken together, the conservation between mammalian hair cells and zebrafish lateral line hair cells makes them a significant tool to investigate disease and damage as well as possible therapeutic interventions.

Forkhead box G1 (Foxg1) is a member of a large family of transcription factors that regulate multiple cellular processes including proliferation, differentiation, and survival (Clark, Halay, Lai, & Burley, 1993). Foxg1 has been implicated during development to increase proliferation of progenitor cells in multiple tissues including the inner ear tissue (Ding, Meng, Kong, He, & Chai, 2020; Hwang, Simeone, Lai, & Wu, 2009; Wong et al., 2019). Recent work shows that Foxg1 is involved in hair cell development and homeostasis in mammalian models (He et al., 2020; Hwang et al., 2009; Zhang et al., 2020). The loss of Foxg1 function results in morphological deformities of the cochlea and sensory cristae as well as altered hair cell polarity and total hair cell complement (Hwang et al., 2009; Pauley, Lai, & Fritzsch, 2006; Zhang et al., 2020). Foxg1 is also implicated in age related hair cell homeostasis through regulation of reactive oxygen species and autophagy (He et al., 2020). Foxg1 interacts directly with critical hair cell development pathways such as canonical Wnt, FGF, and Notch signaling (Akol, Gather, & Vogel, 2022; Ding et al., 2020). The possible functions of Foxg1 have not been studied in the developing zebrafish lateral line or in the context of hair cell regeneration. We seek to uncover the function of Foxg1 in the development and regeneration of the zebrafish lateral line.

Our current study investigates the function of Foxg1a using the *foxg1a^a266^*(Thyme et al., 2019) mutant line to examine development and regeneration of hair cells and support cells in the zebrafish posterior lateral line (pLL). We show that loss of Foxg1a function results in slower pLL primordium migration during the development of the pLL and delayed neuromast formation. We found that significantly fewer hair cells form in the nascent pLL and there is a reduction in proliferating cells in the developing neuromasts. Following regeneration, we found a significant reduction proliferating support cells, hair cells numbers, and Isl1-positive central support cells in *foxg1a^a266^* mutant neuromasts. This work suggests that Foxg1a functions in the neuromast to enable appropriate proliferation and differentiation of cells in the zebrafish lateral line.

## Results

### *foxg1a* is expressed in the developing and regenerating posterior lateral line

Foxg1 function in the development of mammalian inner ear led us to investigate its possible role in the zebrafish lateral line (Ding et al., 2020; Pauley et al., 2006). Work by others has suggested that *foxg1a* is expressed in lateral line neuromasts but as of yet, no functional role has been elucidated (Baek et al., 2022; Lush et al., 2019). We first sought to determine if *foxg1a* is expressed in the developing lateral line. Using whole mount RNA in situ hybridization (WISH) in wild-type embryos we show *foxg1a* is expressed in the migrating pLL primordium at 28 hours post fertilization (hpf) (Fig. 1A). This migrating primordium deposits clusters of cells in its wake that will continue to proliferate and differentiate to give rise to neuromasts containing mechanosensory hair cells and surrounding support cells (Brignull et al., 2009; Ma & Raible, 2009; Thomas et al., 2015). We see expression of *foxg1a* in newly deposited wild-type zebrafish neuromasts at 2 days post fertilization (dpf) (Fig. 1B) and in maturing neuromasts at 5dpf (Fig. 1C). *foxg1a* expression is maintained in the regenerating neuromast at 3 hours post hair cell ablation with the ototoxic aminoglycoside antibiotic neomycin (Fig. 1D; NEO), 1 day post NEO-exposure (Fig.1E), and after regeneration is complete at 3 days post NEO-exposure (Fig. 1F). The continued expression of *foxg1a* in the neuromast during regeneration was also reported by single-cell RNA-sequencing (Baek et al., 2022). These data demonstrate that *foxg1a* expression occurs within the developing and regenerating lateral line.

**Figure 1.**
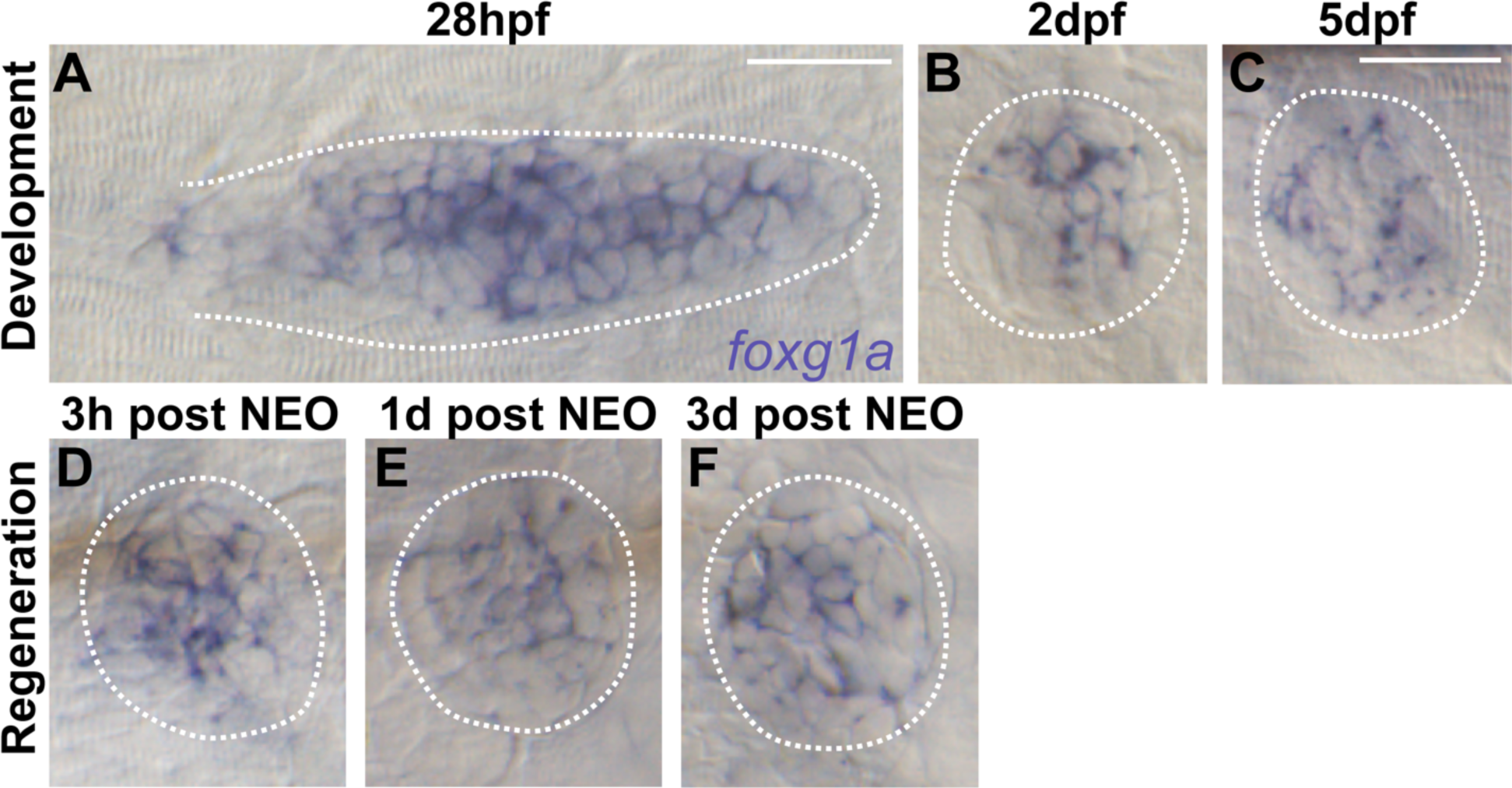
RNA in situ hybridization shows foxg1a expression in developing and regenerating posterior lateral line tissue. (A-C) Whole mount RNA in situ hybridization of *foxg1a* in wild-type zebrafish posterior lateral line primordium at 28hpf (A), and in neuromasts at 2dpf (B) and 5dpf (C). (D-E) Whole mount RNA in situ hybridization of *foxg1a* in wild-type zebrafish neuromasts during regeneration following neomycin (NEO) exposure at 5dpf. (D) 3 hours post-NEO, (E) 1-day post-NEO, and (F) 3 days post. Scale bar = 20μm.

### Foxg1a regulates posterior lateral line primordium migration and neuromast development

As we observed expression of *foxg1a* in the developing and regenerating lateral line, we sought to determine if it plays a functional role as well. During embryonic development, the zebrafish pLL follows a well characterized and stereotypical developmental pattern which involves the collective migration of the pLL primordium cells and deposition of proto-neuromasts along the trunk (Ma & Raible, 2009; Thomas et al., 2015). Mutations affecting pLL development result in phenotypes including truncated lateral line formation, supernumerary neuromasts, loss of neuromasts, altered hair cell numbers, and changes in neuromast size (Lush & Piotrowski, 2014; Thomas et al., 2015). Neuromasts of the zebrafish pLL form after being deposited in the wake of the migrating pLL primordium between ∼22hpf and 48hpf. Live time-lapse imaging of heterozygous and homozygous *foxg1a^a266^* embryos expressing the *Tg(prim:lyn2mCherry)* transgene (Wang et al., 2018) shows pLL primordia migration velocity between 33-47hpf is significantly slower in mutants as compared to controls (Fig. 2A-E, Vid 1). We confirmed this reduced velocity was not due to a reduction in total primordium cells in *foxg1a^a266^* larvae (Fig. S1B,B’’,C) or reduction of cellular proliferation as observed by BrdU incorporation during migration (Fig. S1B,B’,C). At 2dpf we found there was a small, but significant reduction in the number of deposited neuromasts in *foxg1a^a266^* mutants as compared to heterozygous controls (Fig. 2F-H). However, this reduction was recovered by 5dpf (Fig. 2K). These data demonstrate Foxg1a function plays a role in the timing of lateral line development during primordium migration and neuromast deposition.

**Figure 2.**
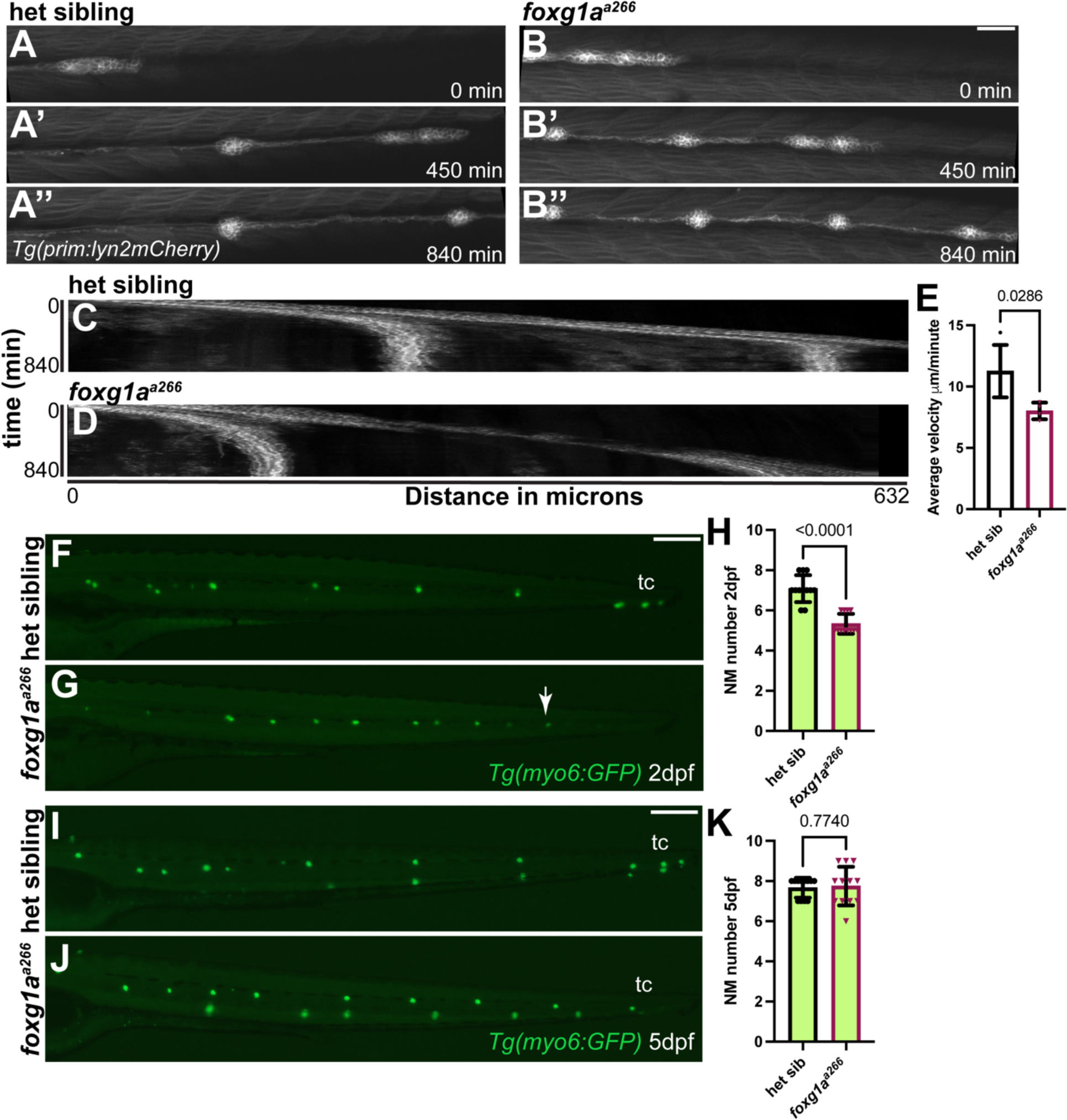
Loss of Foxg1a results in slower posterior lateral line primordium migration and delayed neuromast formation. (A-B’’) Confocal projections of time lapse video of posterior lateral line migration primordium at 0, 450, and 840 minutes in heterozygous (A-A’’) and *foxg1a^a266^* mutant embryos (B-B’’). Scale bar = 50μm. (C-D) Kymograph of time lapse video of posterior lateral line migration in heterozygous (C) and *foxg1a^a266^*mutant embryos (D). (E) Quantification of average primordium velocity during migration, n = 4 embryos per condition. Data presented as mean ±SD. Mann-Whitney U test. (F-G, I-J) Live images of *Tg(myo6:GFP*)-labeled neuromasts in heterozygous (F) and *foxg1a^a266^* mutant embryos (G) at 2dpf and 5dpf in heterozygous (I) and f *foxg1a^a266^* mutant larvae (J). Quantification of NM number at 2dpf (H), n=12 embryos per condition and 5dpf (K), n=12 larvae per condition. Data presented as mean ±SD, Mann-Whitney U test. Scale bars = 20μm.

### Hair cell development is reduced in *foxg1a^a266^* mutant neuromasts

Cells in the maturing zebrafish neruomasts form mechanosensory hair cells and surrounding support cells (Coombs et al., 2014; Ghysen & Dambly-Chaudiere, 2007; Thomas et al., 2015). As Foxg1a seems necessary for proper spatiotemporal migration and neuromast deposition, we next asked if it was necessary for proper hair cell development. Using the *Tg(mysoin6b:GFP)^w186^ (myo6:GFP)* transgenic zebrafish line to label hair cells, we find on average significantly fewer hair cells in *foxg1a^a266^* mutant pLL neuromasts compared to heterozygous siblings (Fig. 3A,A’,B,B’,C). The reduction in hair cells is also seen at 8dpf, suggesting it is not merely developmental delay (Fig. S2A-C) and it is not due to an increase in apoptosis as measured by TUNEL labeling at both 4dpf, and 8dpf (Fig. S3A-H). Hair cells in pLL neuromasts are innervated with afferent and efferent neurons with some mutations that affect lateral line development or damage causing disruption of innervation (Kindt & Sheets, 2018).We also quantified the population of hair cells with functional mechanoelectrical transduction (MET) channels using the fixable fluorescent vital dye FM1-43FX, which enters hair cells through MET channels (Fettiplace, 2017). We observe there is also a significant reduction in FM1-43FX labeled hair cells in *foxg1a^a266^* larvae (Fig. 3A,A’’,B,B’’,C). We next asked if loss of Foxg1a function alters inneration of pLL hair cells using the *Tg(neuroD:eGFP)* transgenic zebrafish line (Obholzer et al., 2008). At 5dpf we do not see a significant change in proportion of hair cells with encircling axons when comparing *foxg1a^a266^*larvae to heterozygous siblings (Fig. 3D,D’’,E,E’’,G). The hair cells of the primary neuromasts in pLL are polarized in an anteroposterior manner and this polarity can be disrupted by mutations in signaling pathways, such as the planar cell polarity pathway (Navajas Acedo et al., 2019). We examined hair cell polarity using actin staining with phalloidin and found no significant difference in hair cell orientation of *foxg1a^a266^* larvae (Fig. 3I,J) compared with heterozygous controls (Fig. 3H,J). These data suggest that the development of neuromast hair cells appear morphologically and functionally intact in *foxg1a^a266^*mutant larvae, though their overall numbers are significantly reduced. However, we must note that these experiments do not fully confirm functional transduction of signals between hair cells and innervating neurons.

**Figure 3.**
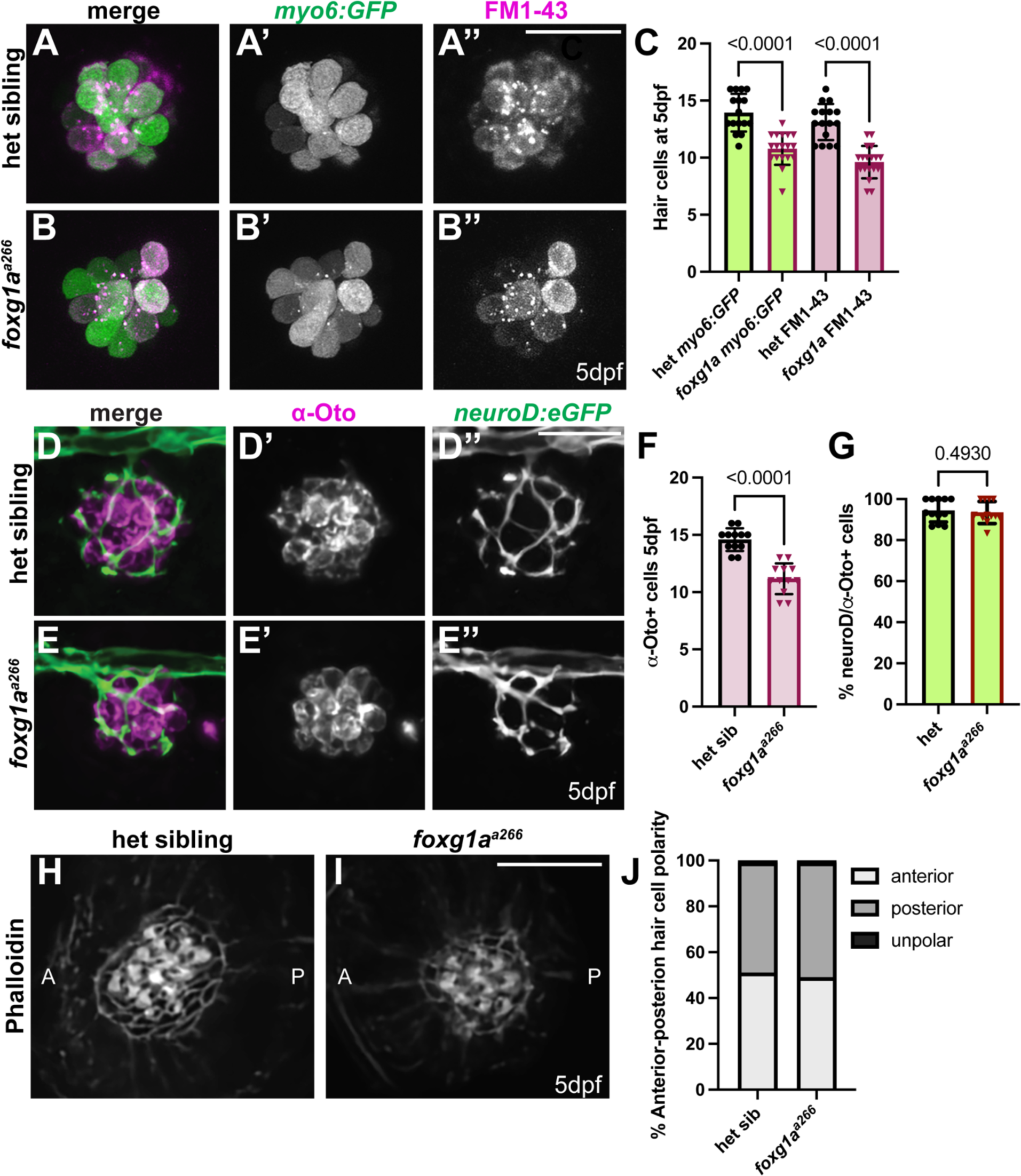
Loss of Foxg1a reduces neuromast hair cell number without loss of innervation or changes in hair cell orientation. (A-B’’) Live confocal projections of *Tg(myo6:GFP)-*labeled (green) and FM1-43FX-labeled (magenta) neuromasts in heterozygous (A-A’’) and *foxg1a^a266^* mutant larvae (B-B’’) at 5dpf. Scale bar = 20μm. (C) Quantification of *myo6(GFP*)+ and FM1-43FX+ hair cells at 5dpf. n= 17 neuromasts in 11 heterozygous sibling larvae, n=19 neuromasts in 10 *foxg1a^a266^*larvae. Data presented as mean ± SD. Kruskal Wallis test with Tukey post hoc comparisons. (D-E’’) Confocal projections of 5dpf heterozygous sibling (D-D’’) and *foxg1a^a266^* (E-E’’) showing hair cells labeled with α-Otoferlin (magenta), and axons expressing *TgBAC(neurod:EGFP*) (green). (F) Quantification of average number of α-Otoferlin+ hair cells per neuromast and (G) quantification of hair cells surrounded by *TgBAC(neurod:EGFP)+* neurons. n=12 neuromasts, 6 larvae per condition. Data presented as mean ±SD, Mann-Whitney U test. Scale bars = 20μm. (H-I) Confocal projects of phalloidin labeled cuticular plates in heterozygous sibling (H) and *foxg1a^a266^* mutant (I) neuromasts. (F) Quantification of the percent anterior-posterior polarized or unpolarized hair cells at 5dpf. n=9 neuromasts in 7 heterozygous sibling larvae and n=12 neuromasts in 8 *foxg1a^a266^* larvae. There is no significant difference in hair cell polarity, Fisher’s exact test. Scale bar=10μm.

### Proliferation is reduced in *foxg1a^a266^* mutant neuromasts

Since we observed a reduction in hair cells without an apparent increase in cell death, we reasoned that the hair cells may not be generated due to effects on cellular proliferation in *foxg1a^a266^* mutant larvae. To investigate this, we used BrdU incorporation in 24-hour pulses between 2-5dpf, during the initial maturation of pLL neuromasts. We find significantly fewer cells demonstrating BrdU incorporation and fewer hair cells in *foxg1a^a266^* mutant neuromasts as compared to heterozygous controls in larvae exposed to BrdU between 2-3dpf (Fig. 4A-A’’,B-B’’,G,H) and between 3-4dpf (Fig. 4D,D’’,J,K). We also find there is a significant increase in the percent of BrdU-labeled hair cells in *foxg1a^a266^* mutant fish pulsed with BrdU from 3-4dpf compared to heterozygous siblings (Fig. 4L). When we observed BrdU incorporation from 4-5dpf, we see that *foxg1a^a266^* larvae show similar BrdU incorporation as compared to heterozygous siblings while still showing a reduction in hair cells (Fig. 4E-E’’,F-F’’,M,N). This data suggests that loss of Foxg1a function results in fewer hair cells as a consequence of reduced proliferation during neuromast development. It also argues that reduced proliferation is not merely the result of developmental delay, as BrdU incorporation is not significantly different between *foxg1a^a266^* mutants and control larvae by 5dpf.

**Figure 4.**
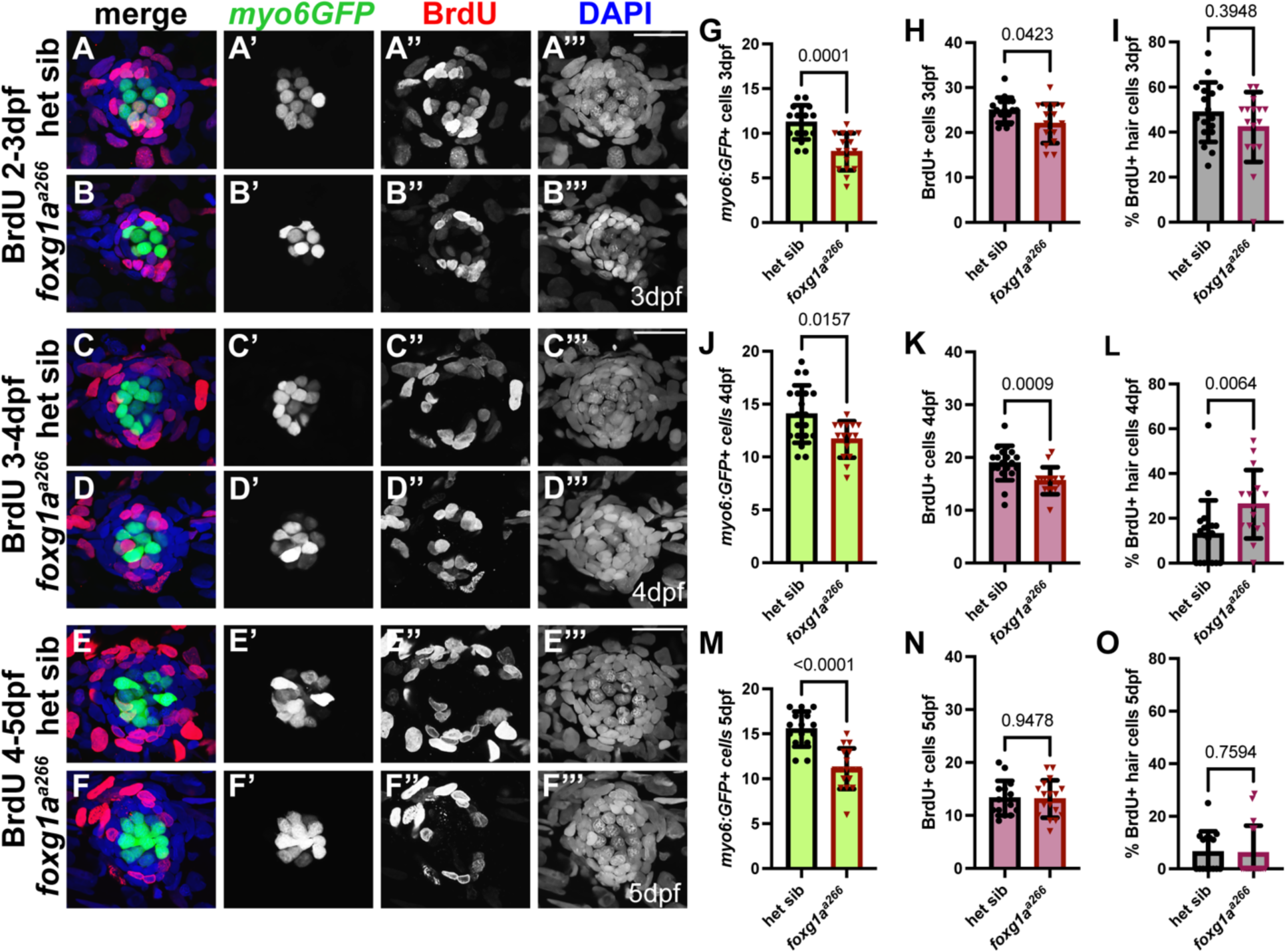
BrdU incorporation in foxg1a mutants is reduced during neuromast maturation. (A-F’’’) Confocal projections of heterozygous sibling and foxg1aa266 embryos expressing *Tg(myosin6B;GFP*) (green) following 24 hour windows of BrdU (red) exposure and nuclei labeled with DAPI (blue) between 2-5dpf. (A-A’’’) 3 dpf heterozygous sibling and (B-B’’’) *foxg1a^a266^* mutant neuromast exposed to BrdU from 2-3dpf), 4 dpf (C-C’’’) heterozygous sibling and (D-D’’’) *foxg1a^a266^*mutant neuromast exposed to BrdU from 3-4dpf), and 5 dpf (E-E’’’) heterozygous sibling and (F-D’’’) *foxg1a^a266^* mutant neuromast exposed to BrdU from 4-5dpf. (G-I) Quantification of heterozygous sibling and *foxg1a^a266^*hair cells (G), total BrdU incorporation (H), and index of BrdU to hair cells (I) at 3dpf. n=17 neuromasts in 9 heterozygous sibling larvae, n=16 neuromasts in 8 foxg1a larvae. (J-L) Quantification of heterozygous siblings and *foxg1a^a266^*hair cells (J), total BrdU incorporation (K), and index of BrdU to hair cells (L) at 4dpf. n=19 neuromasts in heterozygous sibling larvae, n=15 neuromasts in 8 *foxg1a^a266^* larvae. (M-O) Quantification of heterozygous siblings and *foxg1a^a266^* hair cells (M), total BrdU incorporation (N), and index of BrdU to hair cells (O) at 5dpf. n=15 neuromasts in 8 heterozygous sibling larvae. n=17 neuromasts in *foxg1a^a266^* larvae. Mann Data presented as mean ± SD, Mann-Whitney U tests for each condition and time point. Scale bar = 20μm.

### Support cell populations are differentially affected in *foxg1a^a266^*mutant NMs

During development, cells deposited in maturing neuromasts will give rise to hair cells, surrounding support cells, and quiescent mantle cells (Brignull et al., 2009; Ma & Raible, 2009). As loss of Foxg1a function results in fewer hair cells, we asked whether there was also a reduction in the non-sensory support cells within the developing neuromast. To look at total support cell populations we used an α-Sox2 antibody and noted no significant difference in total support cells when comparing heterozygous sibling larvae to *foxg1a^a266^* mutants at 5dpf (Fig. 5A,A’’,B,B’’,D). We also noted there was no change in total DAPI-labeled cells in *foxg1a^a266^* mutants as compared to heterozygous controls (Fig. 5A,A’’’,B,B’’’E), though we observe a reduction in α-Otoferlin (α-Oto)-labeled hair cells (Fig. 5A,A’,B,B’,C). Work by others has demonstrated there are subpopulations of neuromast support cells which show differential function as well as genetic expression (Baek et al., 2022; Lush et al., 2019; Megerson et al., 2024; Thomas & Raible, 2019). Using WISH, we compared the expression profiles of several genes which are expressed in different populations of neuromast cells in heterozygous and *foxg1a^a266^*larvae at 5dpf; these genes include *foxg1a, atoh1a, notch3, deltaD, sost, tnfsf10l3, lfng, and isl1a* (Fig. S4 A-H’), Within this set of genes we find decreased expression of *foxg1a* and *isl1a* in *foxg1a^a266^* mutant neuromasts and compared to heterozygous controls (Fig. S4 A-A’, H-H’)These subpopulations are distinguished based on their transcriptomic profile and spatial organization within the NM (Fig. 5F) (Lush et al., 2019; Thomas & Raible, 2019). We examined the sub-populations using *Tg(sfrp1a:nlsEos)^w217^ (sfrp1a:nlsEos)* to label mantle cells (Thomas & Raible, 2019) (Fig. 5G-H) and *Tg(sost:nlsEos)^w215^*(*sost:nlsEos*) to label dorsoventral cells (Thomas & Raible, 2019) (Fig. 5J-K), and found no significant difference in the number of labeled cells in *foxg1a^a266^* fish as compared to heterozygous siblings (Fig. 5I,L). Using hybridization chain reaction (HCR) fluorescent in situ hybridization (FISH) we quantified the number of cells expressing an anteroposterior support cell marker *tnfsf10l3* at 5dpf and found no significant difference in the number of labeled cells between *foxg1a^a266^* and heterozygous sibling larvae (Fig. 5M-O). Finally, to observe a more basally located central support cell population (Fig. 6A), which along with hair cells is labeled by α-Islet1 (α-Isl1) antibody. We find significantly fewer α-Isl1-labeled cells in *foxg1a^a266^* mutant NMs and heterozygous siblings at 5dpf (Fig. 6B,B’’,C,C’’,E). These data suggest that while Foxg1a is necessary for proper development of hair cells, it does not appear to be required for appropriate differentiation and specification of most support cell populations in pLL neuromasts. Only α-Isl1-labeled hair cells and central support cells show a significant reduction with loss of Foxg1a.

**Figure 5.**
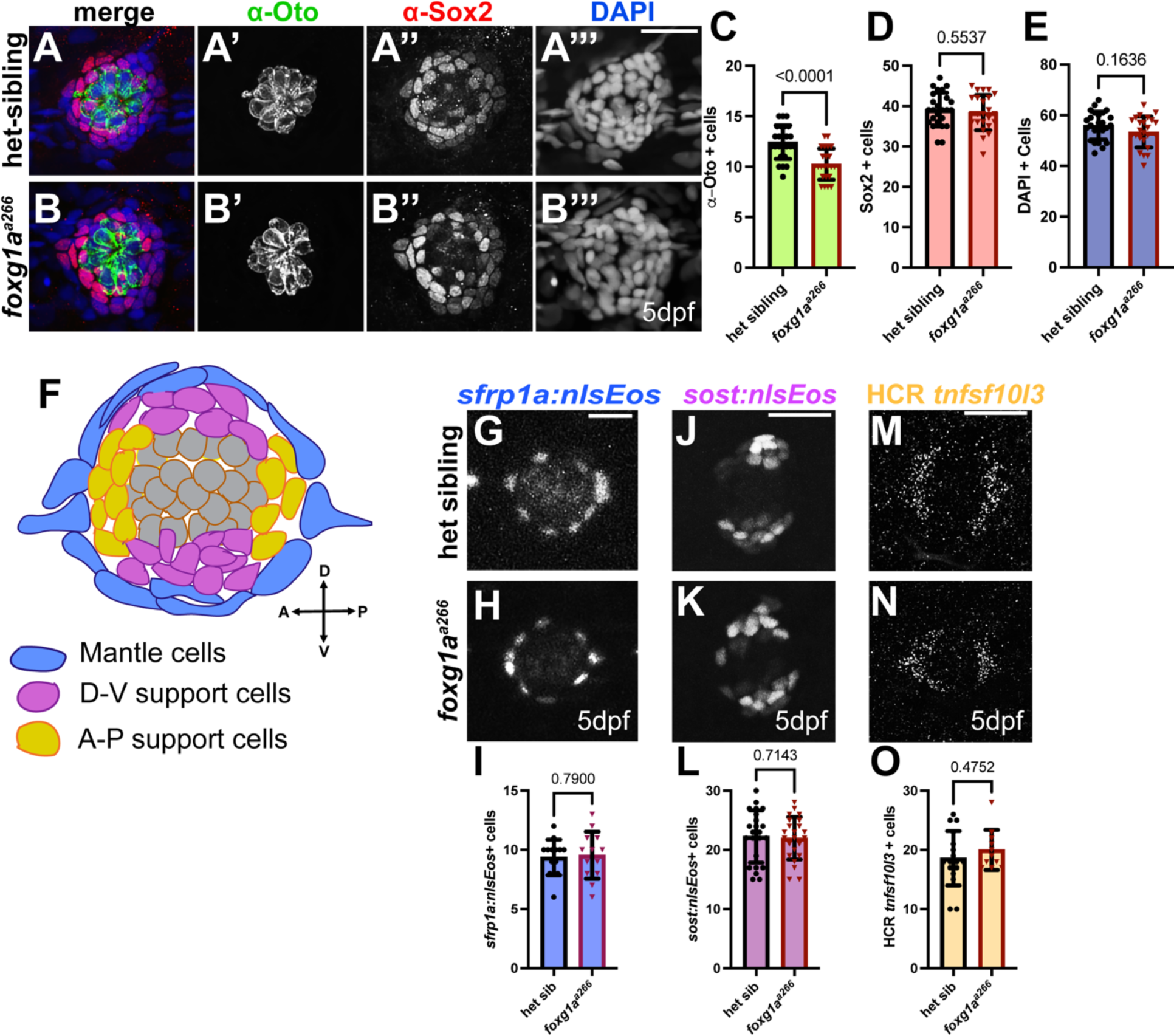
Support cell populations are unaffected by loss of Foxg1a during development. (A-B’’’) Confocal projections of 5dpf (A-A’’’) heterozygous sibling and (B-B’’’) *foxg1a^a266^* larvae labeled with a-Otoferlin antibody-labeled hair cells (green), α-Sox2 antibody-labeled support cells (red), and DAPI-labeled nuclei (blue). (C-E) Quantification of hair cells (C), support cells (D), and total neuromast cells (E) 5dpf. n=25 neuromasts in11 heterozygous sibling larvae, n=24 neuromasts in12 *foxg1a^a266^*. (F) Schematic of support cell populations in zebrafish posterior lateral line neuromast. (G-H) Confocal projection of *Tg(sfrp1a:nlsEos)-*expressing surrounding support cells in 5dpf heterozygous sibling (G) and *foxg1a^a266^*larvae (H). (I) Quantification of *Tg(sfrp1a:nlsEos)-*positive dorsoventral support cells. n=14 neuromasts in 8 heterozygous sibling larvae and n= 13 neuromasts in 6 *foxg1a^a266^* larvae. (J-K) Confocal projection of *Tg(sost:nlsEos*)-expressing dorsoventral support cells in 5dpf heterozygous sibling (J) and *foxg1a^a266^*larvae (K). (L) Quantification of *Tg(sost:nlsEos)-* positive dorsoventral support cells. n=23 neuromasts in 8 larvae per condition. (J-K) Confocal projections of hybridization chain reaction (HCR) fluorescent in situ for *tnfsf10l3* in 5dpf heterozygous sibling (J) and *foxg1a^a266^*larvae (K). (L) Quantification of *tnfsf10l3*+ cells. n= 17 neuromasts in 8 heterozygous larvae and n= 11 neuromasts in *foxg1a^a266^* mutant larvae. All quantification data presented as mean ± SD. Mann-Whitney U test. Scale bar = 20μm.

**Figure 6.**
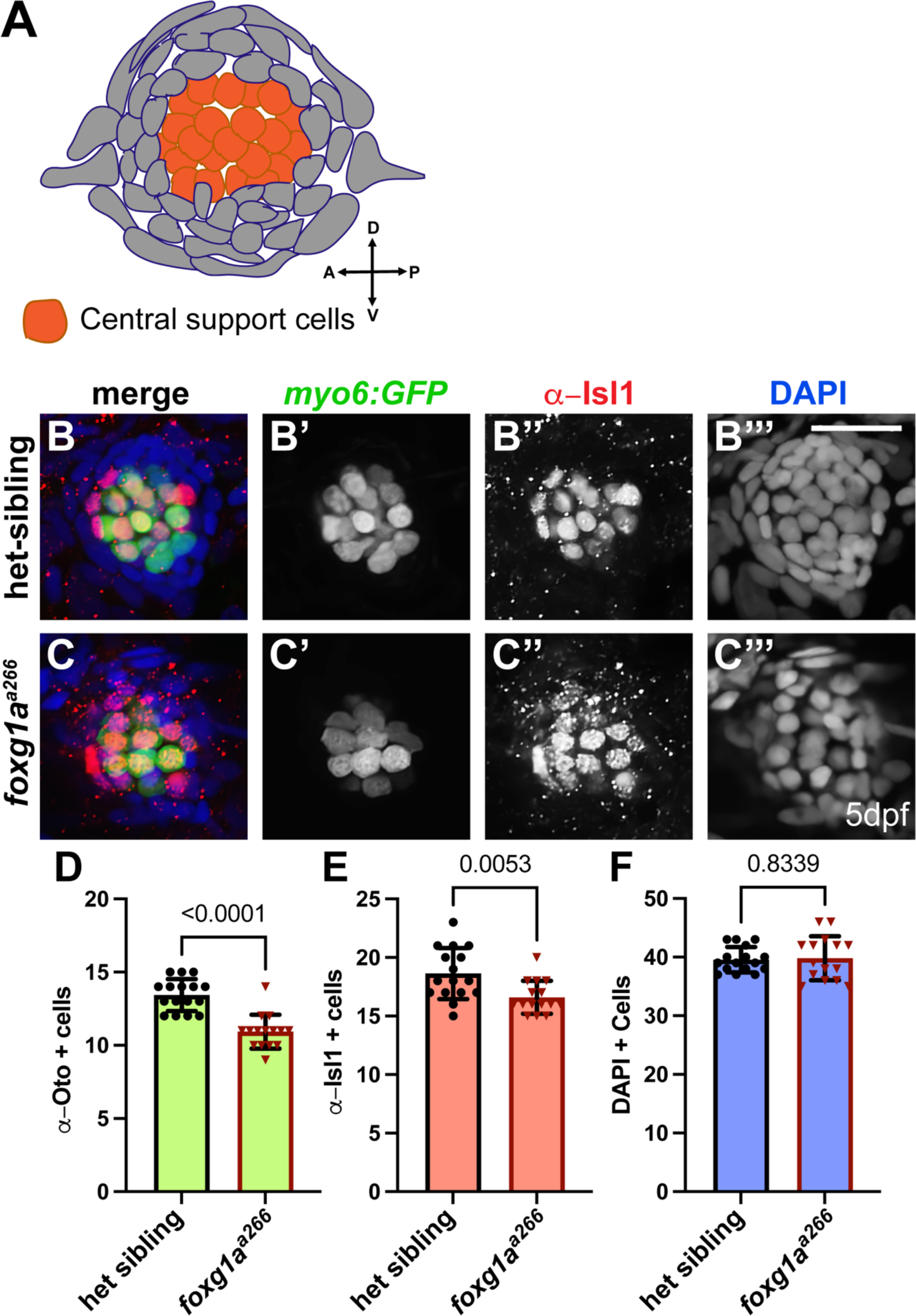
α-Isl1-labeled cells are reduced in *foxg1a^a266^* mutant neuromasts. (A) Schematic of α-Isl1-labeled central support cells and hair cells in the neuromast. (B-C’’’) Confocal projections of 5dpf larvae hair cells expressing *Tg(myosin6B:GFP*) (green), α-Isl1 antibody labeled cells (red), and DAPI-labeled nuclei (blue) in heterozygous sibling (B-B’’’) and *foxg1a^a266^*mutant (C-C’’’) larvae. (D-F) Quantification of hair cells (D), α-Isl1-labeled cells (E), and total neuromast cells (F) at 5dpf.. n= 25 neuromasts in 11 heterozygous sibling larvae and n=24 neuromasts in 12 *foxg1a^a266^* mutant larvae. All data presented as mean ± SD. Mann-Whitney U tests. Scale bar = 20μm.

### Foxg1a is required for hair cell regeneration

Many of the molecular and cellular mechanisms driving hair cell development in the zebrafish lateral line are also active during regeneration (Cruz et al., 2015; Kniss et al., 2016; Pinto-Teixeira et al., 2015). For that reason, we asked if Foxg1a function is required for regeneration of hair cells. We used NEO-exposure to ablate hair cells in 5dpf zebrafish larvae when the majority of hair cells are mature (Harris et al., 2003) and assessed regeneration at time points up to 8dpf. We used FM1-43FX and α-Oto antibody to label total and MET channel-functional hair cells. NEO treated larvae were fixed at 1, 2, and 3 days-post NEO exposure (6, 7, and 8dpf respectively) to track the time course of hair cell regeneration in heterozygous sibling and *foxg1a^a266^*neuromasts (Fig. 7A). At 1d-post NEO (6dpf), we noted no significant difference in the number of regenerated hair cells, either total or with functional MET channels, in *foxg1a^a266^*mutants as compared to heterozygous siblings (Fig. 7B-B’’,C-C’’,D,E). At 2d-post NEO (7dpf), however, we did observe a significant reduction in hair cells labeled with FM1-43FX and α-Oto following regeneration in *foxg1a^a266^* larvae as compared to heterozygous controls (Fig. 7G-G’’,H-H’’,I,J). This reduction in hair cells of *foxg1a^a266^*mutants compared to heterozygous siblings was also observed at our final time point of 3d-post NEO (8dpf; Fig. 7L-L’’, M-M’’, N,O). We did not find a difference between mutant and control larvae when analyzing total neuromast cell numbers using DAPI labeling at each 24-hour interval (Fig 7F,K,P). A TUNEL assay at 18h-post NEO,when proliferation is at its peak (Ma et al., 2008), showed no significant difference in labeling between heterozygous sibling (Fig. S5A-A’’’,D) and *foxg1a^a266^* mutants (Fig. S5B-B’’’,D), suggesting cell death is not the major cause of the reduction of regenerated hair cells in *foxg1a^a266^* mutant larvae.

**Figure 7.**
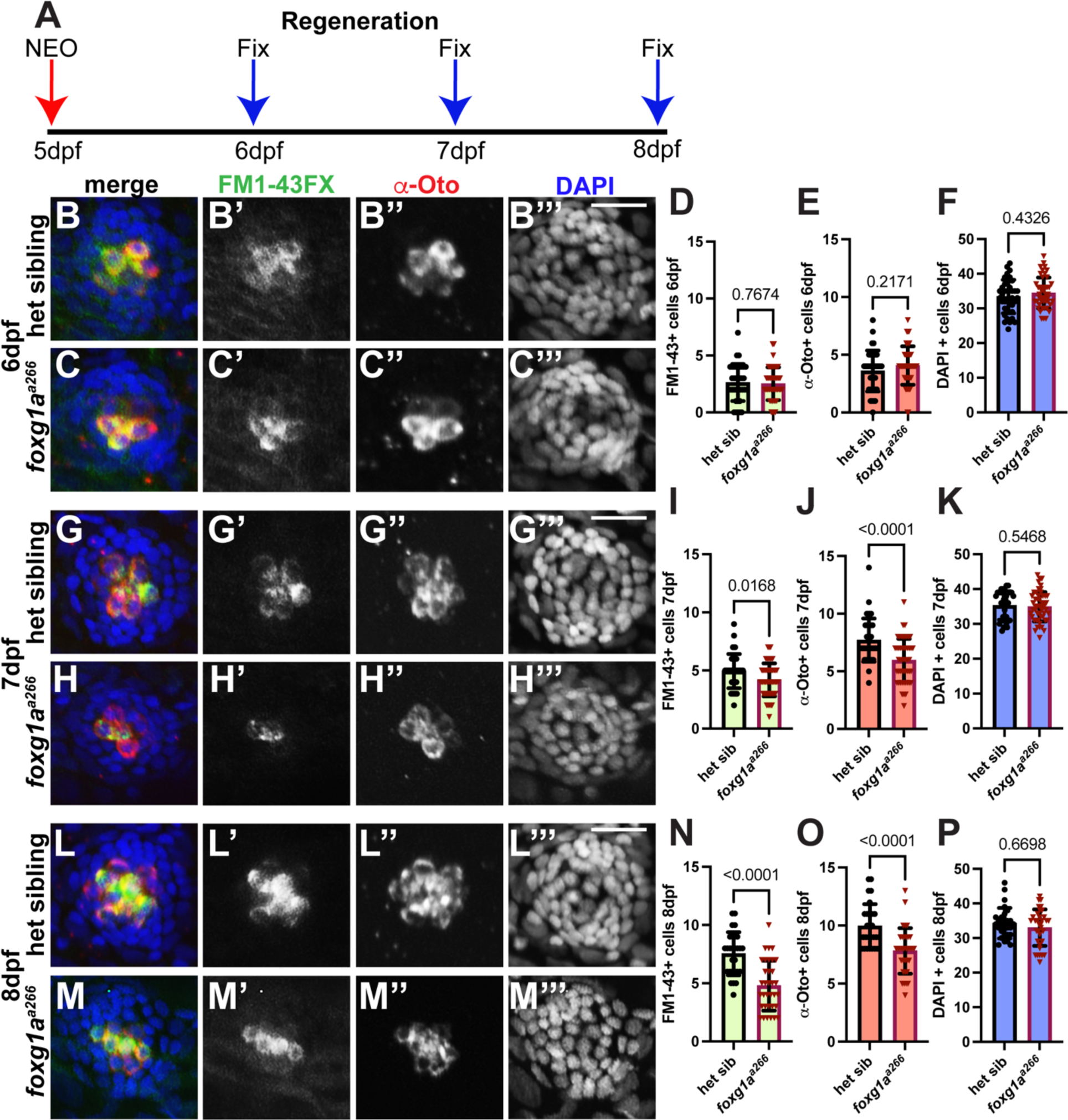
Loss of Foxg1a function results in reduced hair cell regeneration. (A) Schematic of regeneration time line showing fixation at 24 hour increments after NEO-exposure at 5dpf. (B-C’’’, G-H’’’, L-M’’’) Confocal projections of NEO treated zebrafish with hair cells labeled with α-Otoferlin antibody (red) and FM1-43FX (green), and nuclei labeled with DAPI (blue). Regeneration of hair cells at 6dpf in heterozygous sibling (B-B’’’) and *foxg1a^a266^*mutant (C-C’’’) neuromasts. (D-F) Quantification of heterozygous sibling and *foxg1a^a266^*larvae hair cell regeneration and total neuromast cell population 1-day post NEO. n=37 neuromasts in 11 heterozygous and n=46 neuromasts in 12 *foxg1a^a266^* larvae. Regeneration of hair cells at 7dpf in heterozygous sibling (G-G’’’) and *foxg1a^a266^*mutant (H-H’’’) neuromasts. (I-J) Quantification of *foxg1a^a266^*and heterozygous sibling larvae hair cell regeneration and total neuromast cell population 2-days post NEO. n=38 neuromasts in 11 heterozygous sibling larvae and n=51 neuromasts in 10 *foxg1a^a266^* larvae. Regeneration of hair cells at 8dpf in heterozygous sibling (L-L’’’) and *foxg1a^a266^*mutant (M-M’’’) neuromasts. (N-P) Quantification of *foxg1a^a266^*and heterozygous sibling larvae hair cell regeneration and total neuromast cell population 3 days post NEO. n=28 neuromasts in 8 heterozygous sibling and n=34 neuromasts in 8 *foxg1a^a266^* larvae. All data presented as mean ± SD. All conditions analyzed using Mann-Whitney U tests. Scale bars=20μm.

### Proliferation is reduced during regeneration in *foxg1a^a266^* neuromasts

As loss of Foxg1a function results in reduced proliferation in developing neuromasts, we asked if there is a similar reduction during regeneration. To determine if reduced proliferation is a cause of fewer hair cells regenerating in *foxg1a^a266^*mutants, we incubated larvae in BrdU for 24 hours immediately following hair cell ablation with NEO and were then allowed a further two days of regeneration before fixation and analysis (Fig. 8A). We observed a significant decrease in BrdU-labeled cells in the *foxg1a^a266^* neuromasts as compared to heterozygous siblings (Fig. 8B,B’’,C,C’’,E). We next sought to determine whether BrdU-labeling was localized to regenerating hair cells, support cells, or both. We found the percentage of BrdU-labeled hair cells was not significantly different between *foxg1a^a266^*mutant NMs and heterozygous controls (Fig. 8B-B’’, C-C’’,G), indicating that proliferating support cells gave rise to regenerated hair cells. In contrast, when we compare the proportion of BrdU-labeled hair cells to total BrdU-labeled cells in the NM, we found that there was a significant bias toward the hair cell population in *foxg1a^a266^* mutants as compared to controls (Fig. 8B-B’’’, C-C’’’,H), suggesting that proliferating supports cells are preferentially differentiating into hair cells in mutant larvae, perhaps at the expense of self-renewal.

**Figure 8.**
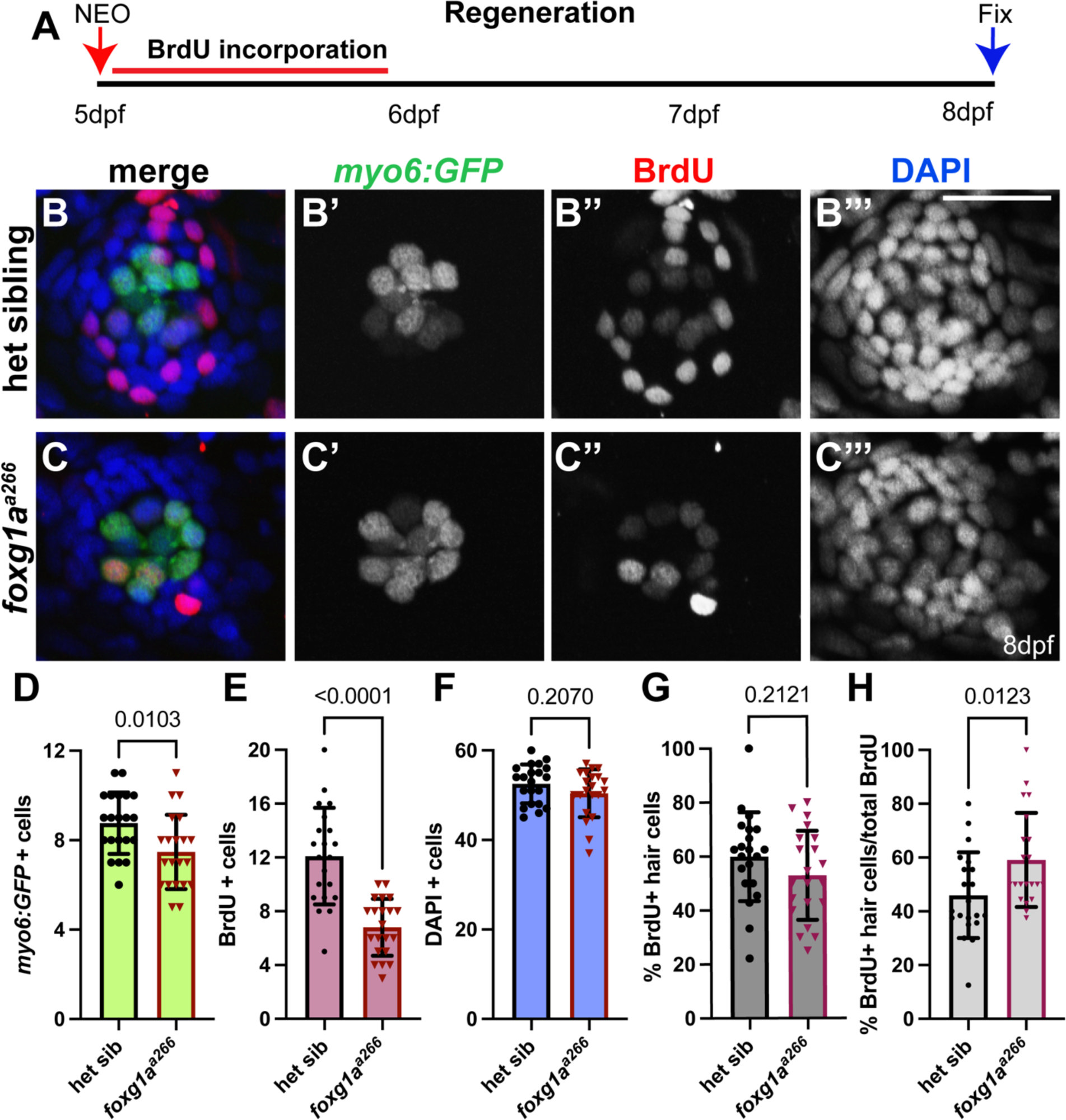
Proliferation is reduced in foxg1aa266 mutants during regeneration. (A) Diagram of of NEO expose at 5dpf, followed by 24 h of BrdU incubation and then regeneration through to 8dpf and fixation. (B-C’’’) Confocal projections at 3 days-post NEO treatment at 8dpf; hair cells and labeled with *Tg(myosin6B:GFP)* (green), proliferating cells are labeled by BrdU-incorporation (red), and nuclei labeled with DAPI (blue) in heterozygous sibling (B-B’’’) and *foxg1a^a266^* mutant (C-C’’’) neuromasts. (D-H) Quantification of hair cells (D), BrdU-labeled cells (E), total nuclei (F), % of BrdU+ hair cells (G), and the %BrdU+ hair cells compared to total BrdU+ cells (H). n=21 neuromasts in 7 larvae per condition. Data presented at mean ± SD. Mann-Whitney U tests. Scale bar = 20μm.

### Foxg1a is required to maintain central support cells following regeneration

During regeneration, differing support cell subpopulations give rise to new hair cells and/or replenish support cells based on location within the zebrafish neuromast (Lush et al., 2019; Thomas & Raible, 2019). Work by others demonstrated that regenerating hair cells arise predominantly from dorsoventral support cells (Romero-Carvajal et al., 2015; Thomas & Raible, 2019) and tend to be located more central to the neuromast as opposed to the periphery (Lush et al., 2019).To track the contribution of dorsoventral cells to hair cell regeneration, we photoconverted *sost:nlsEos-*positive cells prior to NEO-exposure and traced their fate following regeneration (Fig. 9A). We found no significant difference in the total number of *sost:nlsEos* labeled support cells following regeneration comparing heterozygous and *foxg1a^a266^* larvae (Fig. 9B,B’’,C,C’’,E). We also noted no significant difference in the number of *sost:nlsEos-*positive hair cells between *foxg1a^a266^* and heterozygous sibling larvae (Fig. 9B-B’’,C-C’’,D,E,G) This suggests that the *sost;nlsEos*-expressing dorsoventral support cells appear to be contributing to hair cell renewal appropriately in the *foxg1a^a266^*mutants.

**Figure 9.**
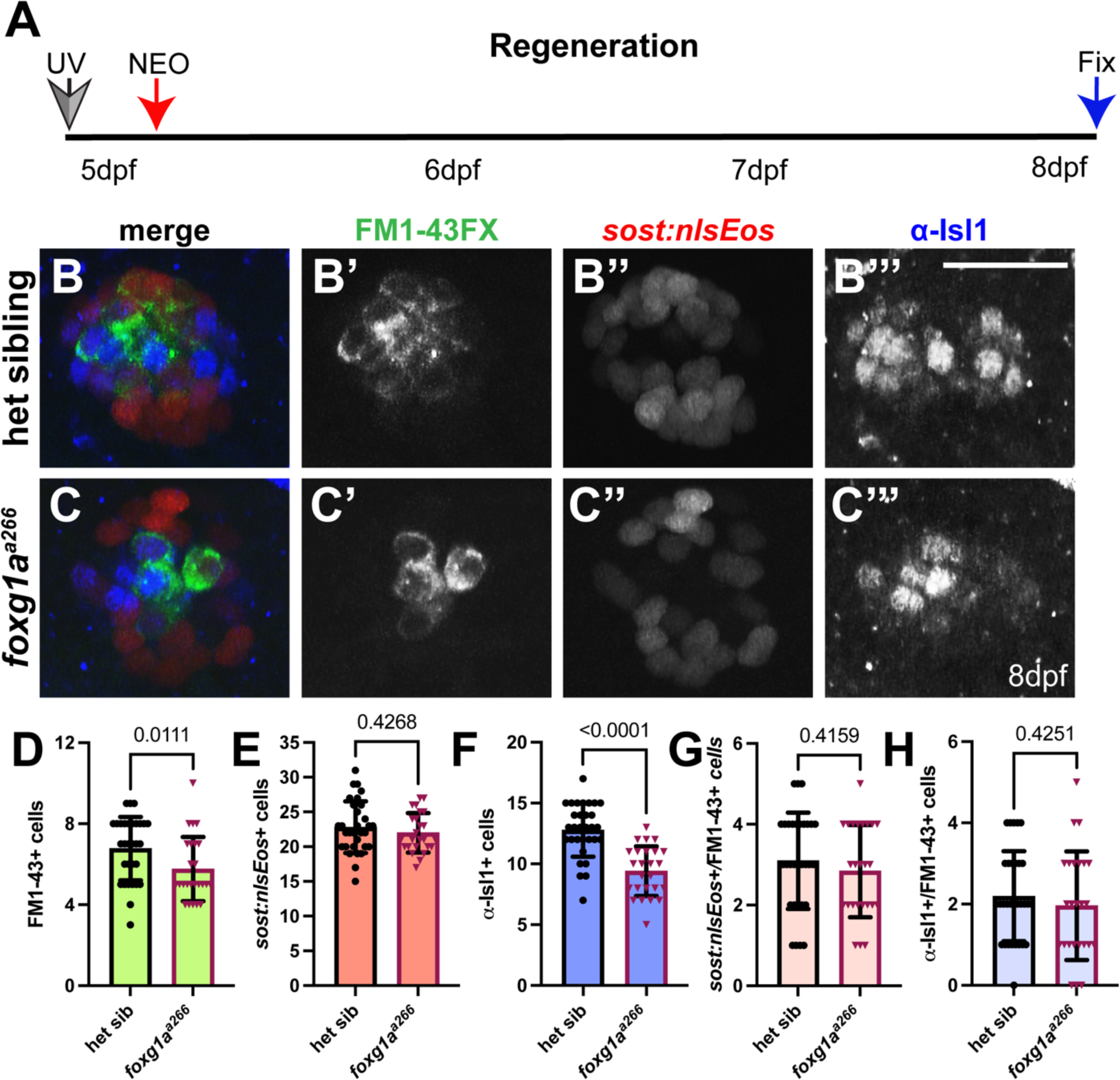
Support cells populations are differentially altered during regeneration in foxg1aa266 mutants. (A) Schematic of UV photoconversion and NEO-exposure prior regeneration. (B-C’’’) Confocal projections of larvae at 8dpf post regeneration showing hair cells labeled with FM1-43FX (green), *sost:nlsEos* dorsoventral support cells (red), and central support cells and hair cells labeled with α-Isl1 antibody (blue). (B-B’’’) heterozygous sibling and (C-C’’’) *foxg1a^a266^* larvae 3d-Post NEO treatment at 8dpf. (D) Quantification of hair cells heterozygous sibling and *foxg1a^a266^* larvae 3d-post NEO. (E) Quantification of *Tg(sost:nlsEos)* labeled hair cells in heterozygous sibling and *foxg1a^a266^* larvae 3d-post NEO. (F) Quantification of α-Isl1-labeled cells in heterozygous siblings and *foxg1a^a266^* larvae 3d-post NEO. (G) Quantification of *sost:nlsEos*+ hair cells in heterozygous siblings and foxg1aa266 larvae 3d-post NEO. (H) Quantification of α-Isl1+ hair cells in heterozygous siblings and *foxg1a^a266^*larvae 3d-post NEO n=31 neuromasts in12 heterozygous sibling larvae and n=25 neuromasts in 11 *foxg1a^a266^* larvae. All data presented at mean ± SD. Mann-Whitney U test. Scale bar = 20μm.

Genetic analysis using scRNA-sequencing suggests that the central support cell populations can also give rise hair cells during regeneration (Lush et al., 2019). As we noted fewer α-Isl1-labeled cells during development in *foxg1a^a266^*mutant larvae, we next asked if these cells showed significant changes following regeneration. Following regeneration, we observed a significant decrease in the number of α-Isl1-labeled cells in *foxg1a^a266^* larvae (Fig. 9C,C’’,F) as compared to heterozygous controls (Fig. 9B,B’’’,F). Most intriguingly, when we compared the number of FM1-43FX-positive hair cells co-labeled with α-Isl1 (Fig. 9B-B’, B’’’, C-C’, C’’’, H) we noted no significant difference between *foxg1a^a266^* mutant larvae and heterozygous siblings, though the number of FM1-43FX-labeled hairs cells was significantly reduced (Fig, 9D), These data suggest that Foxg1a function in the lateral line functions to maintain α-Isl1-positive central support cells and hair cell regeneration.

## Discussion

Foxg1 is a transcription factor that has largely been studied for its role in neural development and subsequent developmental defects accompanying mutations within its gene (Wong et al., 2019; Zhao, Suh, Prat, Ellingsen, & Fjose, 2009). *foxg1a* has been investigated in zebrafish neural development, but our study is the first to look specifically at lateral line development and regeneration (Raj et al., 2020; Toresson, Martinez-Barbera, Bardsley, Caubit, & Krauss, 1998; Zhao et al., 2009). Though recent research suggests *foxg1* is required for the proper development of electroreceptors in the lateral line of paddle fish (Minarik et al., 2024). Our work demonstrates that *foxg1a* is necessary for the proper development and regeneration of the zebrafish pLL. Loss of Foxg1a function results in delayed posterior lateral line development, a reduction in proliferation, and fewer hair cells. During regeneration, *foxg1a* is required for proliferation, formation of new hair cells, and the maintenance of Isl1-postive central support cells in the neuromast. We note that α-Isl1 antibodies labels hair cells in *foxg1a^a266^* mutants at similar numbers to controls and the precise role of the Isl1-postive central cells during regeneration remains unknown.

### Foxg1a functions to promote proper development of mechanosensory systems

In studies of the mammalian inner ear, Foxg1 is demonstrated to be necessary for proper morphological formation, innervation, and sensory cell development (Ding et al., 2020; Hwang et al., 2009). Foxg1 knockdown in mice embryos results in shortened cochlea, greater proportion of inner ear hair cells, and a reduction in inner ear hair cell innervation (Ding et al., 2020; Hwang et al., 2009; Pauley et al., 2006; Zhang et al., 2020). When we looked at early pLL development in *foxg1a^a266^* mutant zebrafish, we noted a reduced rate of migration of the posterior lateral line primordium with recovery by 5dpf and deposition of terminal cluster neuromasts. It is interesting to note that murine models investigating the knockout of Foxg1 demonstrated an increase in hair cells (Pauley et al., 2006), in contrast to the zebrafish lateral line where we see a decrease in hair cell development. In mice, the loss or conditional knockdown of Foxg1 results in significant disruption to innervation of hair cells and the cochlea (Pauley et al., 2006; Zhang et al., 2020). When we examined innervation of hair cell in the *foxg1a^a266^*, we found a similar pattern to controls, suggesting there axonal extension of the sensory neurons associated with pLL may not be regulated by Foxg1a. More work is needed to better distinguish the precise role of Foxg1 mechanosensory system development as there is evident conservation of function within the tissue though possibly differing effects between species.

### Foxg1a is necessary for proliferative development and regeneration of sensory tissue

Hair and support cells of the zebrafish lateral line develop and regenerate mostly through proliferation and differentiation of support cell progenitors (Coombs et al., 2014; Ghysen & Dambly-Chaudiere, 2007; Kniss et al., 2016; Ma & Raible, 2009; Thomas et al., 2015). As mammalian Foxg1 is implicated in proliferative development of nervous tissue we reasoned the zebrafish Foxg1a may also regulate proliferation (Akol et al., 2022; Wong et al., 2019). Our work shows that loss of Foxg1a function results in reduced proliferation both during development and regeneration of the zebrafish pLL without an increase in cell death. Interestingly, we note that following NEO treatment, regenerated hair cells still show proportional BrdU incorporation compared to controls, while overall BrdU-labeled cells are decreased, suggesting that support cells are preferentially forming hair cells. Our work suggests a conserved role for Foxg1in support cell entry into the cell cycle (Akol et al., 2022; Ding et al., 2020; Hwang et al., 2009). The timing of Foxg1 function may also be important as murine work with loss of function (Pauley et al., 2006) or conditional KD (Zhang et al., 2020) showed differing effects on between different developmental time points. This is also important as regeneration in murine models is observed during a short window in early development while zebrafish hair cells retain the capacity to regenerate through the life of the animal (Brignull et al., 2009; Matsui & Cotanche, 2004).

We noted that loss of Foxg1a function resulted in a decrease in the central support cell subpopulation of the neuromast labeled by an α-Isl1 antibody. The centrally located cell population has been shown to be the predominant locus for proliferation resulting in new hair cells during regeneration (Lush et al., 2019). Additionally, work by others has demonstrated that support cells with a dorsoventral orientation contribute more to hair cell regeneration than anteroposterior or peripheral support cell populations (Thomas & Raible, 2019). When we analyzed *foxg1a^a266^* and heterozygous controls we noted both demonstrated hair cells were labeled with α-Isl1 and *sost:nlsEos* following regeneration. Taken together we believe this argues that hair cells arise proliferatively from a transcriptionally heterogeneous population of support cells during regeneration. Recent work for the McGraw lab has also demonstrated that some neuromast support cells appear poised to replenish hair cells through non-proliferative mechanisms as well as proliferative (Megerson et al., 2024). As *foxg1^a266^* mutants show reduced proliferation in neuromast during development and regeneration, we propose that this reduction in proliferation results in the decrease in the numbers of hair cells and α-Isl1-positive central support cells.

### Islet1 in the posterior lateral line central support cells

Islet1 is a LIM-homeodomain transcription factor known to regulate development of nervous tissue, neural epithelia, and otic tissue (Liang et al., 2011; Sun et al., 2008). In mice its expression is noted in developing support cell populations and nascent hair cells, with its expression in hair cells fading away as differentiation continues (Radde-Gallwitz et al., 2004). Work by others has shown *isl1* is expressed in the central support cells of zebrafish neuromasts (Lush et al., 2019). Our work recapitulates that these *isl1a* expressing central support cells contribute to hair cell renewal during regeneration and additionally that loss of Foxg1a results in a decrease in the numbers of this support cell population. These data indicate there is possibly a role for *isl1a* in support cell progression to a hair cell fate that is at least in part regulated by *foxg1a.* Future studies will address the precise function of Foxg1a in central support cells, in particular the target genes regulated by Foxg1a transcriptional function. It would also be interesting to investigate other known cellular functions of Foxg1a like autophagy, regulation of ROS, and cell cycle.

### Concluding remarks

Our study reveals a novel function of Foxg1a in regulating cellular proliferation, central support cell numbers, and hair cell development and regeneration in the zebrafish posterior lateral line. We believe this work provides greater insight into the cellular and molecular mechanisms driving vertebrate mechanosensory tissue development and may elucidate new opportunities in mammalian hair cell regeneration.

## Methods and Materials

### Zebrafish lines and maintenance

The following *Danio rerio* (zebrafish) lines were used: wild-type*AB (ZIRC;http://zebrafish.org), *foxg1a^a266^* (Thyme et al., 2019), *Tg(sost:nlsEos)^w215^* (Thomas and Riable, 2019), *Tg(myosin6b:GFP)*^w186^ (Thomas and Raible, 2019), *Tg(sfrp1a:nlsEos)^w217^* (Thomas and Raible, 2019),*Tg(prim:lyn2mCherry)* (Wang et al., 2018), *TgBAC(neurod:EGFP)^nl1^*(Obholzer et al., 2008), and *(-4.9sox10:EGFP)^ba2^* (Dutton et al., 2009). Zebrafish were maintained and staged according to standard protocols (Kimmel, Ballard, Kimmel, Ullmann, & Schilling, 1995). Experiments reported in this study were conducted on larvae between 24 hours post fertilization (hpf), and 8 days post fertilization (8dpf). Larvae were kept in E3 embryo medium (14.97 mM NaCl, 500 μM KCL, 42 μM Na_2_HPO_4_, 150 μM KH_2_PO_4_, 1 mM CaCl_2_ dihydrate, 1 mM MgSO_4_, 0.714 mM NaHCO_3_, pH 7.2). For all experiments, larvae were treated with tricaine (Syndel) prior to fixation in 4% paraformaldehyde/PBS (Thermo Fisher). All research was performed in accordance with the McGraw laboratory protocol #45344 approved by the UMKC IACUC committee. In laboratory zebrafish lines, sexual determination and differentiation occurs at ∼25dpf (Kossack & Draper, 2019), after the timepoints used in this study.

## METHOD DETAILS

### Whole mount RNA in situ hybridization

Whole mount RNA in situ hybridization (WISH) was carried out using established protocols (Thisse & Thisse, 2008), modified with a 5-minute Proteinase K (Thermo Fisher) treatment to preserve neuromast integrity. The probes used were: *atoh1a* (Itoh & Chitnis, 2001), *notch3* (Itoh & Chitnis, 2001), *deltaD* (Haddon et al., 1998)*, foxg1a, tnfsf10l3*, *sost, sfrp1a, lfng, and isl1a.* Antisense probes were generated using established protocols (Thisse & Thisse, 2008) or using a PCR-based protocol (Logel, Dill, & Leonard, 1992).

### HCR fluorescent RNA in situ hybridization

Hybridization chain reaction (HCR) fluorescent in situ hybridization (FISH) was carried out following the manufacturer’s protocol. (Molecular Instruments). The probes used were *isl1a*-B1(4 pmol), *atoh1a-*B2(4 pmol) and *tnfsf10l3-*B2 (4 pmol), with the amplifiers B1-546 and B2-647 respectively (Molecular Instruments). Larvae were subsequently labeled with DAPI and mounted using Fluorescent Mounting Media (EMD Millipore) to prevent fading.

### Immunohistochemistry, FM1-43FX and DASPEI labeling

Whole mount immunolabeling was performed using established protocols (Ungos et al., 2003) for all antibodies except when noted. The primary antibodies used were: α-Otoferlin antibody (α-Oto; mouse monoclonal, 1:200, DSHB, University of Iowa), α-Sox 2 antibody (rabbit polyclonal, 1:100, Invitrogen), α-Islet1 antibody (α-Isl1; mouse monoclonal, 1:100, DSHB, University of Iowa), and α-BrdU antibody (mouse monoclonal, 1:100, BD Biosciences). For α-Isl1 antibody labeling, a 1 hour room temperature, or 4 degree Celsius overnight water wash was conducted following fixation for 1-2hours at room temperature in 4%PFA/PBS. Larvae in α-Isl1antibody were incubated for 2 days at room temperature. Secondary antibodies used were: goat α-rabbit Alexa-647 antibody (1:1000, Invitrogen), goat α-mouse Alexa-568 antibody (1:1000, Invitrogen), and goat α-mouse Alexa-647 antibody (1:1000, Invitrogen). Mature hair cells were labeled by a 1-minute incubation in 3mM FM1-43FX (Invitrogen; (Owens et al., 2008)). Nuclei were labeled with 30mM DAPI (Thermo Fisher). Antibody block was made per protocol using 2% goat serum. Hair cells were visualized in live larvae using 2-(4-(dimethylamino)styryl)-N-ethylpyridinium iodide (DASPEI; Invitrogen) according to established protocols (Harris et al., 2003).

### TUNEL Labeling

TUNEL labeling was conducted using the Click-iT Plus TUNEL Assay Alexa Fluor 594 (Invitrogen, C10618). The kit protocol was adapted for larval zebrafish; larvae were fixed in 4% paraformaldehyde in PBS for 1hr at room temperature. Proteinase K digestion was done with 10ug/mL proteinase K in PBS for 10 minutes. All wash steps were doubled to reduce background.

### Neomycin exposure and regeneration

For hair cell ablation, 5 days post fertilization (dpf) larvae were incubated in 400mM neomycin (NEO, Millipore-Sigma) in embryo medium for 0.5 hours and then washed 3x in fresh embryo medium. In experiments analyzing complete regeneration, larvae were collected 3 days following NEO exposure. Shorter time periods of regeneration we also conducted per experimental need. For TUNEL labeling following neomycin exposure, larvae were collected 1 hour post NEO. For RNA in situ hybridization, larvae were collected 28hpf, 2dpf, and 5dpf; for regeneration, larvae were collected at 3-hours post-NEO, 1-day post-NEO, or 3 days post-NEO.

### BrdU incorporation

Cellular proliferation was analyzed using 5-bromo-2’-deoxyuridine (BrdU; Millipore Sigma). BrdU incorporation was carried out using established protocols (Harris et al., 2003; Laguerre, Soubiran, Ghysen, Konig, & Dambly-Chaudiere, 2005); larvae were incubated in 10 mM BrdU for 24 hours at set times during development or following NEO-induced ablation, larvae were exposed to BrdU immediately after NEO for 24 hours and then transferred to fresh E3 for 2 days and fixed at 8dpf.

### Photoconversion

For photoconversion experiments using *Tg(sost:nlsEos)^w215^*or *Tg(sfrp1a:nlsEos)^w217^* fish, 5 dpf larvae were placed in a shallow depression slide and exposed to 405 nm light for 20 seconds using a Zeiss Imager.D2 compound microscope and a 10x objective. For regeneration experiments, photoconversion was carried out prior to NEO exposure. For live imaging of *Tg(sost:nlsEos)^w215^*fish, larvae were anesthetized using tricaine and embedded in 1.2% low melt agarose/E3 embryo medium.

### Image collection

For imaging of RNA in situ hybridization and immunohistochemistry, processed larvae were placed in 50% glycerol/PBS and mounted on slides. For imaging of HCR in FISH, larvae were mounted on slides in Fluorescent Mounting Media (Calbiochem) and imaged within 3 days of processing to prevent signal loss. Images were collected using a Zeiss 510 Meta LSM confocal microscope with Zen 2009 software or a Nikon AXR resonant scanning confocal microscope with NIS-Elements software. Images were processed using Fiji software (Schindelin et al., 2012) and brightness and contrast were adjusted using Photoshop (Adobe).

## QUANTIFICATION AND STATISTICAL ANALYSIS

For quantification of neuromast cell numbers, we took the mean of individual neuromasts from multiple fish and conducted appropriate analyses comparing control and mutant samples. Neuromasts from L1-L4 were analyzed for each fish and cells were manually counted. All statistical analysis was carried out using GraphPad Prism 10 (GraphPad Prism version 10.0.0 for Mac, GraphPad Software, San Diego, California USA, www.graphpad.com). The data presented represents discrete variables so use of nonparametric tests determined. A Mann-Whitney U two-tailed nonparametric test was used for pair-wise comparisons, a Kruskal-Wallis test with Dunn’s multiple comparison tests between multiple conditions, and a Fisher’s exact test was used to compare condition across groups. Significance was set at p<0.05. All data is presented as ± standard deviation (SD). A power analysis was conducted from initial samples of hair cell averages with an alpha of 0.05, and beta of 0.2 which established a minimum sample size of 6 per group for counts. Power calculated via ClinCal.com sample size calculator for two independent samples.

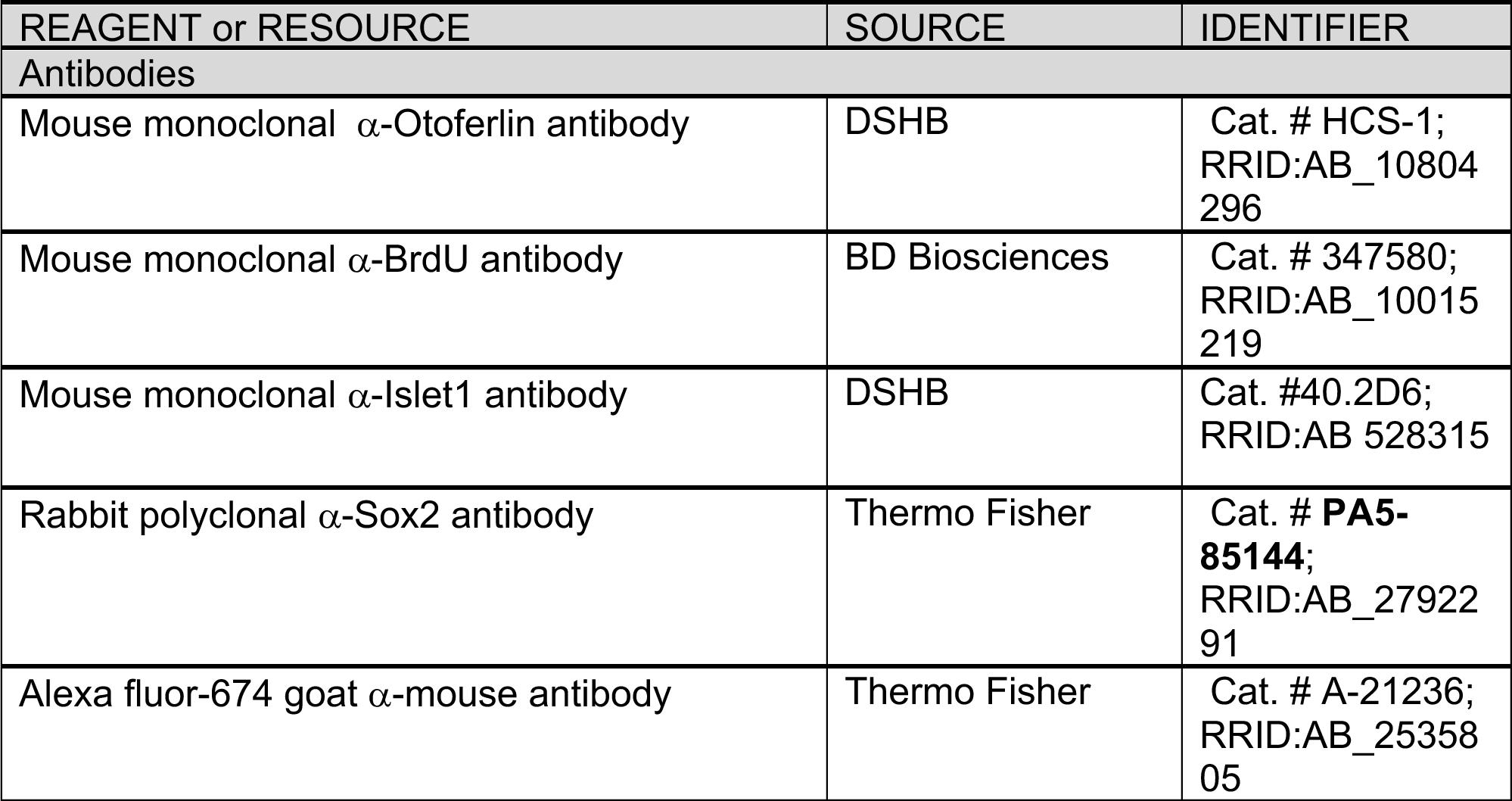
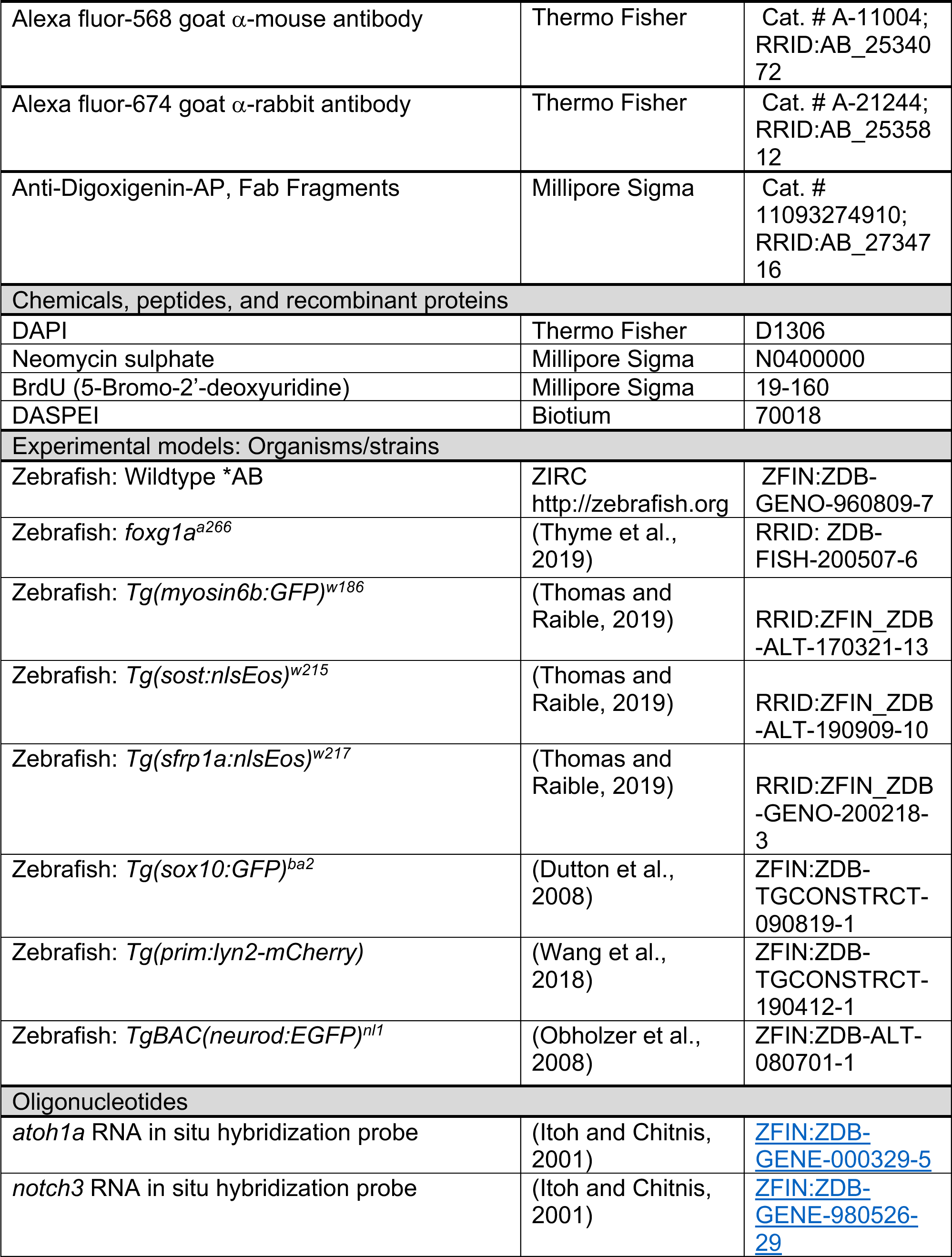
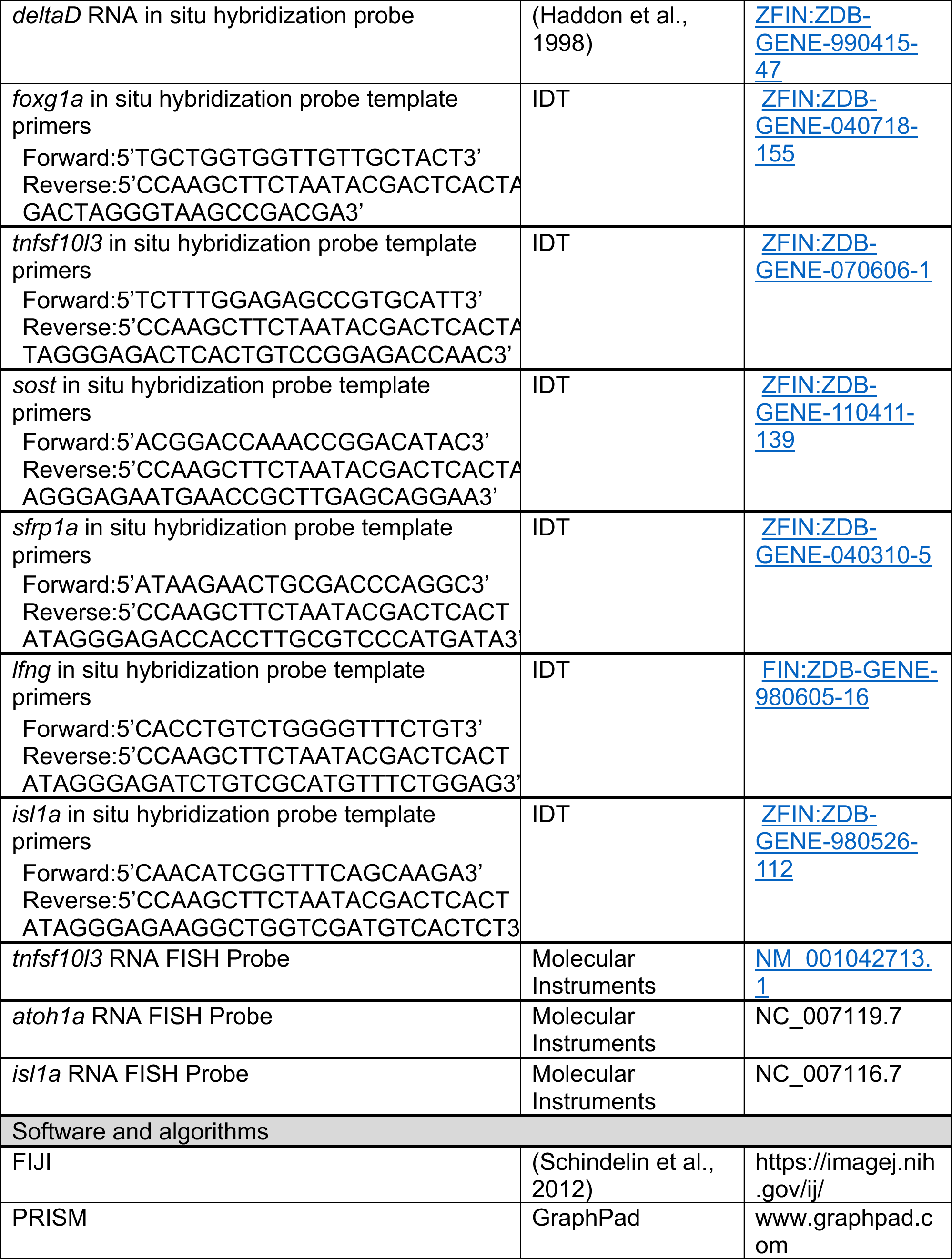

## Acknowledgements

The authors would like to thank Dr. David Raible for providing the, *Tg(myosin6b:GFP)^w186^*, *Tg(sost:nlsEos)^w215^* and *Tg(sfrp1a:nlsEos)^w217^* zebrafish lines. The authors would like to thank Dr. Alexander Schier for the *foxg1a^a266^* zebrafish line. Funding provided by NIGMS 1R16GM146690-01 (HFM).

## Author contributions

Conceived and designed experiments: JMB, CB, EMB, MMV, HFM. Preformed experiments: JMB, CB, EMB, MMV, HFM. Analyzed data: JMB, CB, EMB, MMV, HFM. Wrote Manuscript: JMB, HFM. Edited Manuscript: JMB, HFM.

**The authors declare no competing interests with this work.**

## Notes

### Competing Interest Statement

The authors have declared no competing interest.

