## Supplemental files for "*foxg1a* is required for hair cell development and regeneration in the zebrafish lateral line"

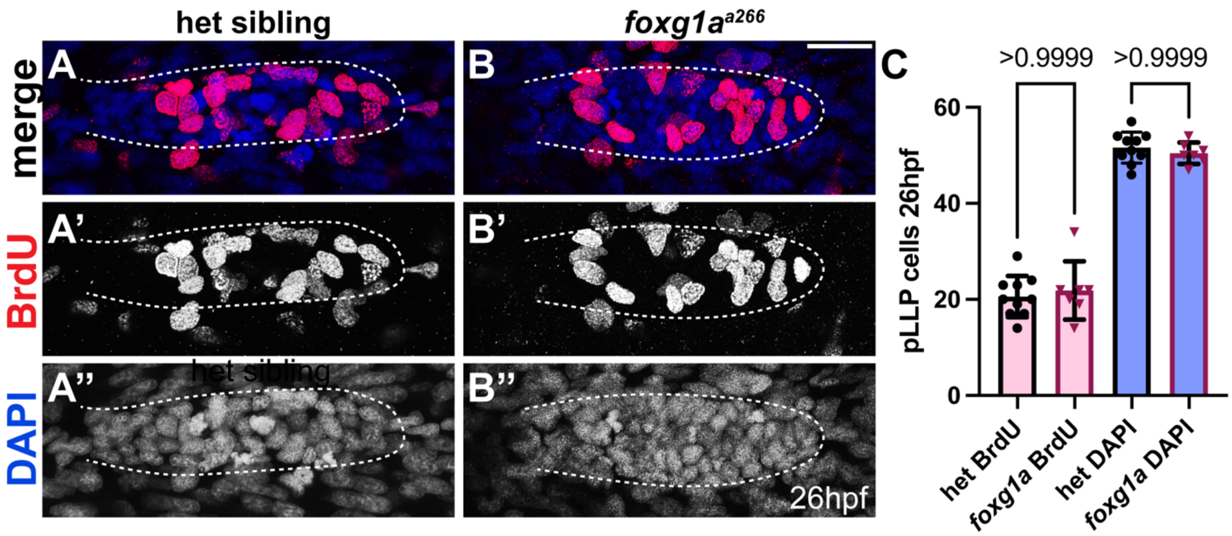


**Supplemental Figure 1. Loss of Foxg1a does not affect proliferation or size of migrating posterior lateral line primordium.** (A-B'') Confocal projections of posterior lateral line primordium at 26hpf. Nuclei labeled with BrdU (red) and DAPI (blue) in heterozygous siblings (A-A'') and *foxg1a*$^{a266}$ (B-B'') embryos. (C) Analysis of total DAPI labeled cells and BrdU labeled cells in migrating primordium 26hpf. n=10 heterozygous sibling, and n=7 *foxg1a.* Data presented as mean ± SD. Kruskal Wallis test with Tukey post hoc comparisons. Scale bar = 20μm.

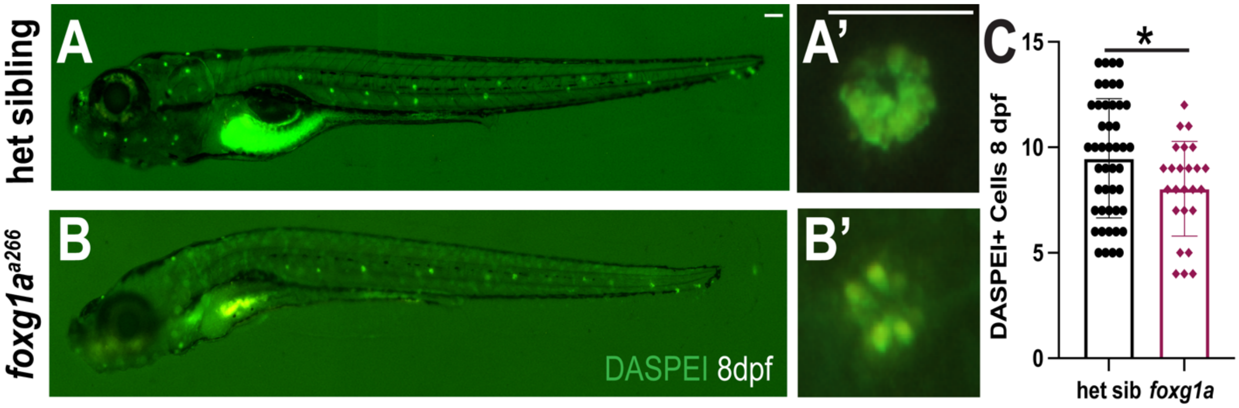


**Supplemental Figure 2. Reduction in hair cells persists through 8 days post fertilization in *foxg1a* mutants.**(A-B') Light microscope projections of lateral line hair cells labeled with DASPEI (green) at 8dpf in heterozygous sibling (A-A') and *foxg1a*$^{a266}$ mutant (B-B') larvae (C) Analysis of DASPEI labeled hair cells 8dpf. All data presented at mean ± SD. Mann-Whitney U test. (A-B) Scale bar=100μm, (A'-B') Scale bar=20μm .

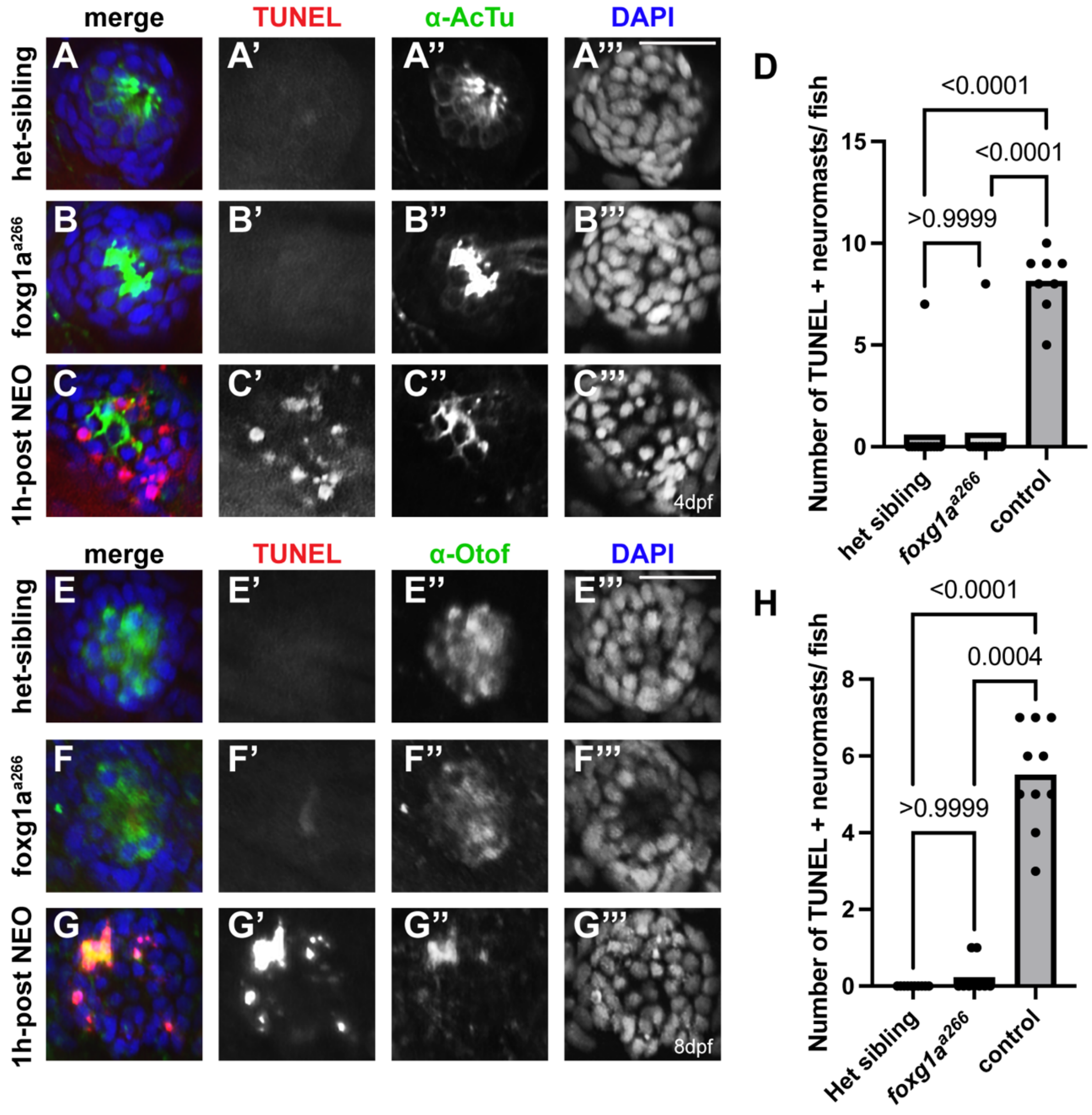


**Supplemental Figure 3. Reduction in hair cell not due to loss from apoptosis.** (A-B''') Confocal projections of heterozygous sibling and *foxg1a*$^{a266}$neuromasts 4dpf. (C-C''') Confocal projections of control heterozygous sibling neuromasts 4dpf, 1 hour-post NEO. Hair cells marked with $\alpha$-Acetylated Tubulin antibody (green), TUNEL-labeling (red) and nuclei labeled with DAPI (blue),. (D) Percent TUNEL positive cells to TUNEL negative cells per condition. n=110 neuromasts in12 *foxg1a*$^{a266}$ heterozygous sibling larvae, and n=121 neuromasts in 12 *foxg1a*$^{a266}$ *mutant* larvae, and n=65 neuromasts in 8 control wild-type larvae. Fisher's exact test. (E-F''') Confocal projections of heterozygous sibling and *foxg1a*$^{a266}$ neuromasts 8dpf. (G-G''') Confocal projections of control heterozygous sibling neuromasts 8dpf, 1 hour-post NEO. Hair cells marked with $\alpha$-Otoferlin antibody (green), TUNEL-labeling (red), and nuclei labeled with DAPI (blue),. (H) Percent TUNEL positive cells to TUNEL negative cells per condition. n=35 neuromasts in 10 heterozygous sibling larvae, and n=35 neuromasts in 9 *foxg1a*$^{a266}$ mutant larvae, and n=63 neuromasts in 10 control larvae,35 neuromasts. Fisher's exact test. Scale bar = 20$\mu$m.

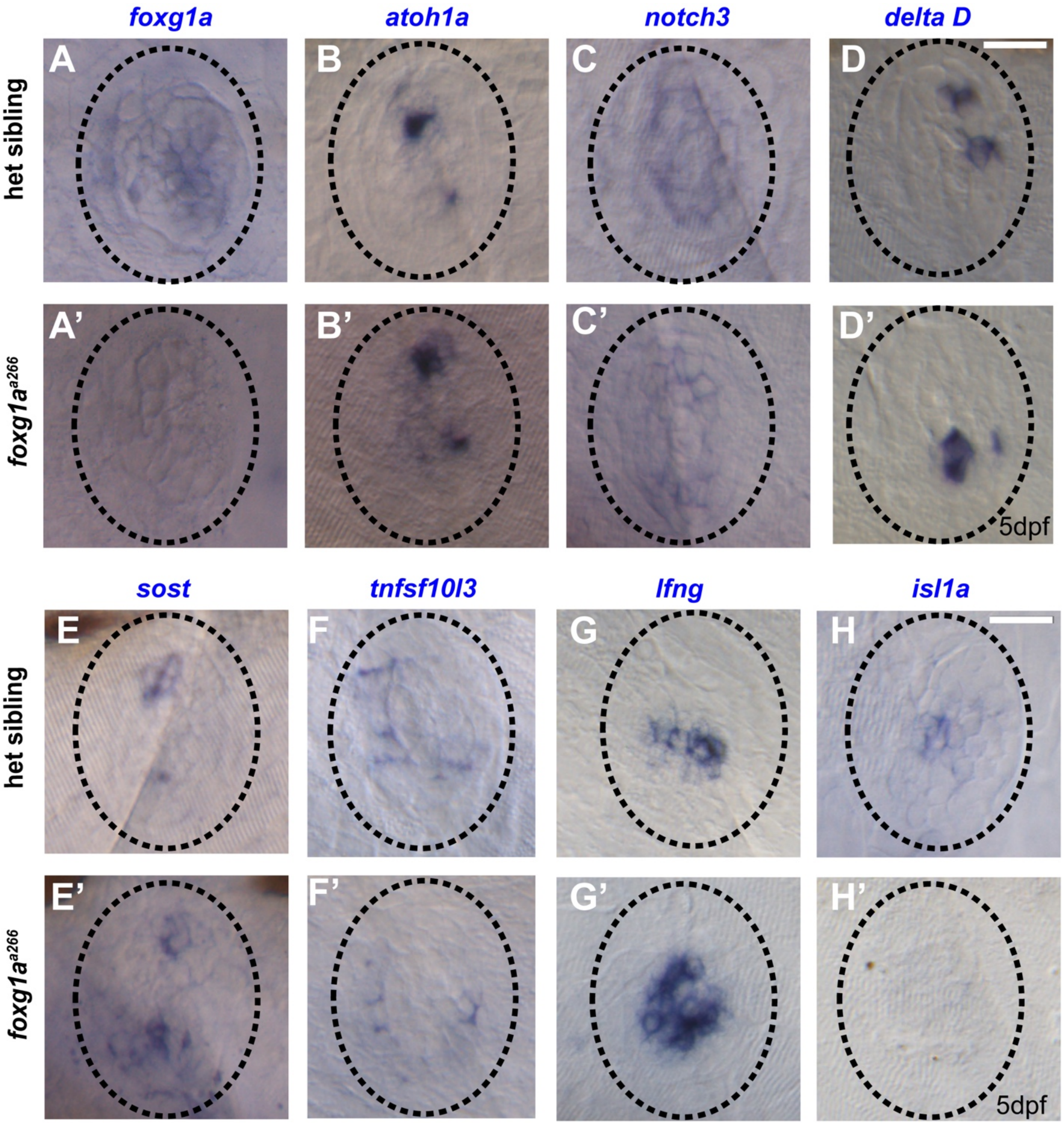

foxg1a
atoh1a
notch3
delta D
A
B
C
D
het sibling
A'
B'
C'
D'
foxg1a^a266
5dpf
sost
tnfsf10l3
lfng
isl1a
E
F
G
H
het sibling
E'
F'
G'
H'
foxg1a^a266
5dpf

**Supplemental Figure 4. RNA in situ hybridization of different cellular markers in *foxg1a* mutants.**(A-A') RNA in situ hybridization showing expression of *foxg1a* in heterozygous sibling and *foxg1a*$^{a266}$ neuromasts 5dpf. (B-B") RNA in situ hybridization showing expression of *atoh1a* in heterozygous sibling and *foxg1a*$^{a266}$ neuromasts 5dpf. (C-C') RNA in situ hybridization showing expression of *notch3* in heterozygous sibling and *foxg1a*$^{a266}$ neuromasts 5dpf. (D-D') RNA in situ hybridization showing expression of *deltaD* in heterozygous sibling and *foxg1a*$^{a266}$ neuromasts 5dpf. (E-E') RNA in situ hybridization showing expression of *sost* in heterozygous sibling and *foxg1a*$^{a266}$ neuromasts 5dpf. (F-F') RNA in situ hybridization showing expression of *tnfsf10l3* in heterozygous sibling and *foxg1a*$^{a266}$ neuromasts 5dpf. (G-G') RNA in situ hybridization showing expression of *lfng* in heterozygous sibling and *foxg1a*$^{a266}$ neuromasts 5dpf. (H-H') RNA in situ hybridization showing expression of *islet1a* in heterozygous sibling and *foxg1a*$^{a266}$ neuromasts 5dpf. Scale bar = 20μm.

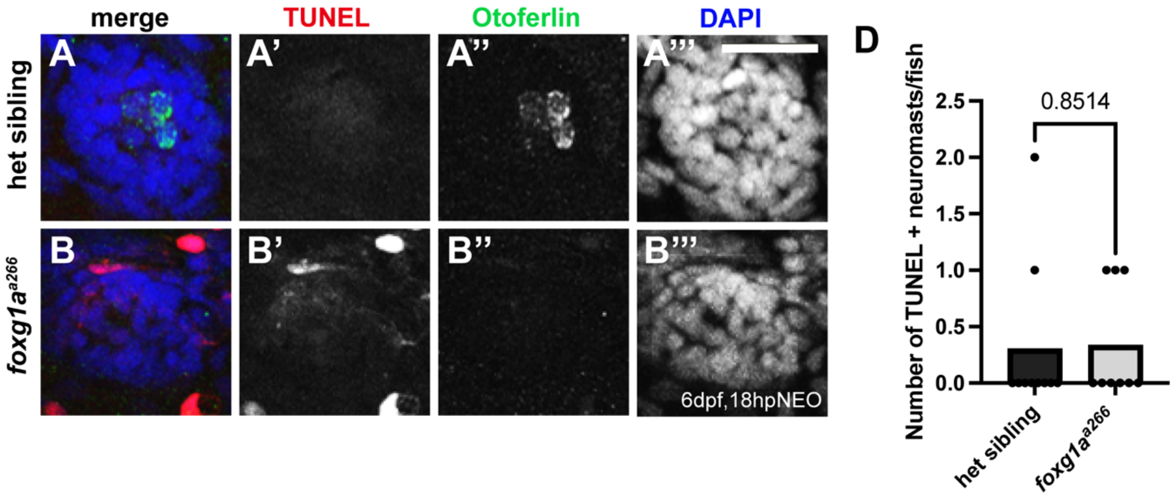


**Supplemental Figure 5. Loss of Foxg1a does not increase cell death during regeneration.** (A-B''') Confocal projections of heterozygous sibling (A-A''') and *foxg1a*$^{a266}$ mutant (B-B''') neuromasts 6dpf, 18 hour-post NEO. Hair cells marked with α-Otoferlin antibody (green), TUNEL-labeling (red), and nuclei labeled with DAPI (blue),. (H) Percent TUNEL-positive cells to TUNEL-negative cells per condition. n=37 neuromasts in12 heterozygous sibling larvae and n=31 neuromasts in 9 *foxg1a*$^{a266}$ larvae. Fisher's exact test. Scale bar = 20μm.
